# Volumetric mesoscopic electrophysiology: a new imaging modality for the non-human primate

**DOI:** 10.1101/2024.05.13.593946

**Authors:** Tobias Teichert, László Papp, Ferenc Vincze, Nioka Burns, Baldwin Goodell, Zabir Ahmed, Andrew Holmes, Charles M Gray, Maysam Chamanzar, Kate Gurnsey

**Affiliations:** Department of Psychiatry, University of Pittsburgh, Pittsburgh, PA; Department of Bioengineering, University of Pittsburgh, Pittsburgh, PA; Neuronelektród Kft, Budaörs, Hungary; Plexon Inc – Neuroscience Technology, Dallas, TX; Graymatter Research, Bozeman, MT; Department of Electrical and Computer Engineering, Carnegie Mellon University, Pittsburgh, PA

## Abstract

The primate brain is a densely interconnected organ whose function is best understood by recording from the entire structure in parallel, rather than parts of it in sequence. However, available methods either have limited temporal resolution (functional magnetic resonance imaging), limited spatial resolution (macroscopic electroencephalography), or a limited field of view (microscopic electrophysiology). To address this need, we developed a volumetric, mesoscopic recording approach (**MePhys**) by tessellating the volume of a monkey hemisphere with 992 electrode contacts that were distributed across 62 chronically implanted multi-electrode shafts. We showcase the scientific promise of MePhys by describing the functional interactions of local field potentials between the more than 300,000 simultaneously recorded pairs of electrodes. We find that a subanesthetic dose of ketamine –believed to mimic certain aspects of psychosis– can create a pronounced state of functional disconnection and prevent the formation of stable large-scale intrinsic states. We conclude that MePhys provides a new and fundamentally distinct window into brain function whose unique profile of strengths and weaknesses complements existing approaches in synergistic ways.

## 1. Background

The primate brain is a highly interconnected organ. Understanding its function requires not only information about sensory input and motor output, but also detailed knowledge of the rich network of functional connections that enables, shapes, and broadcasts intrinsic activity across the entire brain ^1^. This interconnected nature can only be fully appreciated by recording from the entire brain in parallel rather than parts of it in sequence ^2,3^. Hence, the ability to non-invasively record from the entire brain with millimeter spatial resolution is at the core of the appeal of functional magnetic resonance imaging (**fMRI**) ^4–8^. However, the temporal resolution of fMRI is limited by the relatively slow hemodynamic response ^9,10^.

In contrast to fMRI, electrophysiological approaches have excellent temporal resolution. However, they have either 1) limited spatial resolution, as is the case for macroscopic recordings such as scalp electro-encephalography (**EEG**) ^5^, or (2) a limited field of view, as is the case for microscopic single-cell recordings. Despite rapid technical developments ^11–19^, the goal of simultaneously recording from all neurons in a primate brain is not yet on the horizon. Hence, it is critical to develop intermediate electrophysiological approaches that ideally record from across the entire brain, albeit not at the single cell level. To date, such mesoscopic approaches are underdeveloped, thus leaving a valuable source of information mostly untapped.

To address this methodological need, we developed a volumetric, mesoscopic recording approach by tessellating the volume of an entire monkey hemisphere (including cerebrum, cerebellum, diencephalon, and parts of the mesencephalon) with 992 electrode contacts. The contacts were distributed across 62 chronically implanted multi-electrode shafts arranged into 14 coronal slices with 2-to-7 shafts per slice and 16 contacts per shaft. The contacts formed an approximately regular three-dimensional grid with a pitch of 5 mm between shafts in the antero-posterior dimension and 4 mm in the medio-lateral dimension, and less then 3 mm along the shaft.

Note that spacing out electrode contacts in a three-dimensional grid across an entire hemisphere is counterintuitive from a single-cell perspective: by design, many contacts will end up in white matter, and even if it were possible to record individual neurons on each contact, moving the electrodes by as little as 100 micrometers would lead to a completely different set of neurons and results. However, our system was built to record local field potentials (**LFP**), whose spatial profile is smoother^20^, leading to meaningful correlations for distances of up to 10 mm ^21^, thus exceeding the distances between adjacent electrode contacts. LFPs can also aggregate across populations of neurons and form dipole-like structures that propagate through gray and white matter and can reliably be recorded many centimeters from their site of origin in the form of EEG potentials on the scalp ^22–27^. By sampling LFPs in a three-dimensional grid, we will be recording the same type of activity that underlies EEG and ECoG recordings, but with a much finer volumetric spatial grid and higher signal-to-noise ratio. We are referring to this new imaging modality that combines the millisecond temporal resolution of electrophysiology with the large field of view and millimeter spatial resolution of fMRI as **volumetric mesoscopic electrophysiology (MePhys**).

Technically, MePhys is most closely related to stereotactic EEG ^28,29^ (SEEG), i.e., LFP recordings from several chronically implanted multi-contact shafts that are used in the clinic for the localization of seizure foci. MePhys also builds on decades of technical expertise from EEG ^30,31^, and ECoG ^32^ recordings in humans, as well as a wide range of electrophysiolgocal techniques in animal model systems using macroelectrodes ^33–41^, micro-electrodes ^42–44^, linear arrays ^45–48^, eCog arrays ^49,50^, single-tipped electrode-arrays ^3,12,13,17,51,52^, or arrays of multi-channel electrodes ^53^. Conceptually, however, MePhys fundamentally differs from existing approaches along at least one of three dimensions: (i) Its field of view is a three-dimensional volume rather than one- or two-dimensional manifold; (ii) MePhys spreads electrode contacts as widely as possible in order to maximize the field of view rather than spatial resolution; (iii) MePhys prioritizes LFPs over single-cell activity. The conceptual priorities and design choices of MePhys give rise to a unique functional profile that is unlike any existing electrophysiological approach. MePhys has a wide range of potential applications, among them the description of large-scale hemisphere-wide functional networks with high spectro-temporal resolution that is not available in classical fMRI-based functional connectivity analyses.

In the work below we show that MePhys in the monkey is technically feasible, safe, and provides stable hemisphere-wide LFP recordings over a period of months, with no noticeable degradation of signal quality over time. Such hemisphere-wide recordings provide important spatial context that is lacking when recording from only one or few brain regions. We showcase the scientific promise of MePhys by exploring how a subanesthetic dose of ketamine can blunt hemisphere-wide responses to external stimuli, create a pronounced state of functional disconnection and prevent the formation of stable large-scale intrinsic states. We conclude that the unique design and conceptual priorities of MePhys provide a new and distinct window into non-human primate brain function whose profile of strengths and limitations complements existing approaches in synergistic ways. Leveraging its unique functional profile that fills gaps at the intersection of existing methods, MePhys has the potential to facilitate the translation of findings across scales (micro – meso – macro); species (rodent – monkey – human); recording modalities (electrophysiology – fMRI – optical); time-scales (milliseconds – seconds – minutes); and intrinsic states such as motivation, attention or learning (naïve – proficient – overtrained).

## 2. Results

### 2.1 The MePhys System

Designing and implanting a MePhys system with the desired features posed a number of technical challenges. 1) We needed to custom design and manufacture 62 laminar depth probes with variable lengths and inter-electrode spacings as well as a recording stability spanning months to years. 2) It was crucial to design a guide-tube system that would a) allow all 62 electrode shafts to penetrate the dura without buckling or breaking, and b) subsequently keep infections from entering the brain along any of the electrode shafts. 3) Because the large number of electrode shafts could not all be implanted in a single surgery, we needed to design a physical platform that remained (a) accessible to us over the course of successive surgeries, and (b) secure from the monkey. 4) In order to relate the mesoscopic to macroscopic recordings, the system had to accommodate the implantation of EEG electrodes in the skull. 5) Finally, because it takes hours to connect such a large number of electrodes, they needed to remain permanently connected to the head-stage. Hence, the platform needed to provide enough space on the animal’s head for both the head-stage and the wires connecting the stage to the electrodes.

We designed the MePhys platform to solve these challenges by combining three rugged structural elements (*‘crown’*, *‘wall’*, and *‘cap’*), two internal elements that facilitate implantation of the macroscopic and mesoscopic electrodes (*‘spider’* and *‘grid’*), and a 1024 channel head-stage (Spikegadget) and its *‘baseplate’* (**Fig. 1**A & Section 4.2 for details). A computed tomograpy (**CT**) image collected after implanting the MePhys platform (Section 4.4) determined that 58 grid holes were suitable for electrode shaft implantation (Section 4.5). Over the course of the next 4 surgeries, we successfully implanted 57 electrode shafts, 50 of which remained fully functional through all surgeries and to this day, i.e., between 8 and 14 months as of March 2024 (**Fig. 1**B & **Suppl. Fig. 3** & Sections 4.5&4.6). A CT image collected after all surgeries allowed us to visualize trajectories of all electrode shafts, and locations of all 800 functional electrodes (**Fig. 1**C & **Suppl. Fig. 3**).

**Figure 1.**
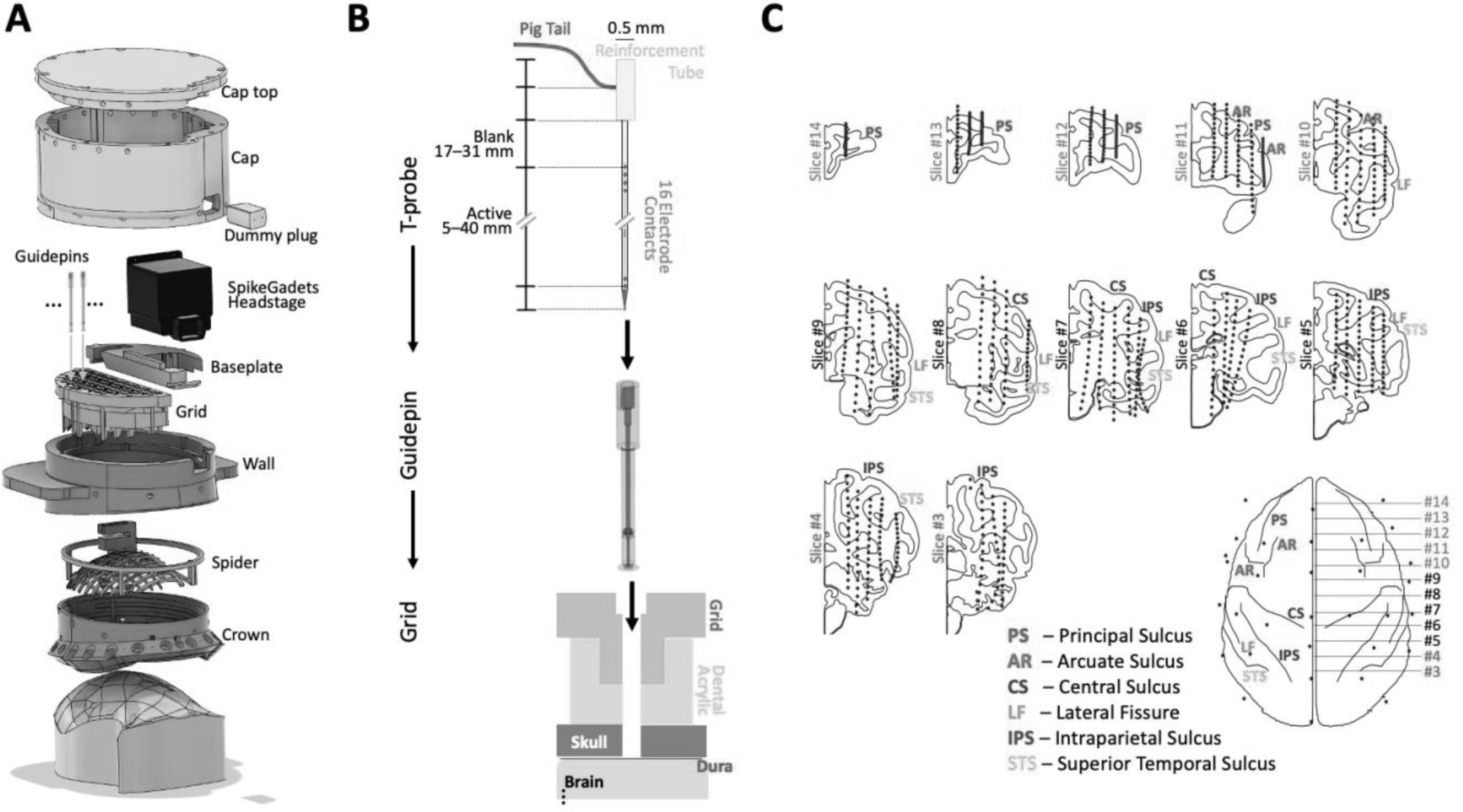
Mesoscopic Electrophysiology (MePhys). (A) Exploded design of the MePhys platform with the main structural and functional elements: crown, spider, wall, grid, baseplate, headstage and cap. (B) Schematic of the electrode – guidepin – grid system. (C) The intracranial contacts (blue dots) of the MePhys prototype are arranged in 14 coronal slices from posterior to anterior in the panels from bottom right to top left. The antero-posterior position of the slices is indicated by horizontal lines in the top-down schematic of the monkey brain. Slices #1 and #2 were omitted, because they do not contain any currently functional electrode contacts.

### 2.2 Sensory Input – Auditory Evoked Electric Fields

#### Baseline stimulus-evoked activity

To showcase the ability of MePhys to track hemisphere-wide stimulus-evoked activity, we recorded electric activity from macro- and mesoscopic electrodes in response to short white-noise bursts of varying intensity at three key time-points of the response (**Fig. 2A-C** & **Suppl. Fig. 4**). (Supplement: animated GIF showcasing propagation of auditory evoked electric fields along the auditory hierarchy). We were able to simultaneously record auditory evoked electric fields from regions in the canonical ascending auditory pathway including brainstem, inferior colliculus, thalamus, and a dense cluster of regions in and around primary auditory cortex (**Fig. 2A-D**). In addition, we also observed auditory evoked responses in the insula; cerebellum; and prefrontal, posterior parietal, and motor cortices. The MePhys data allowed us to quantify evolving functional properties along the ascending auditory pathway such as response latency, duration, and dependence on stimulus amplitude (data not shown).

**Figure 2.**
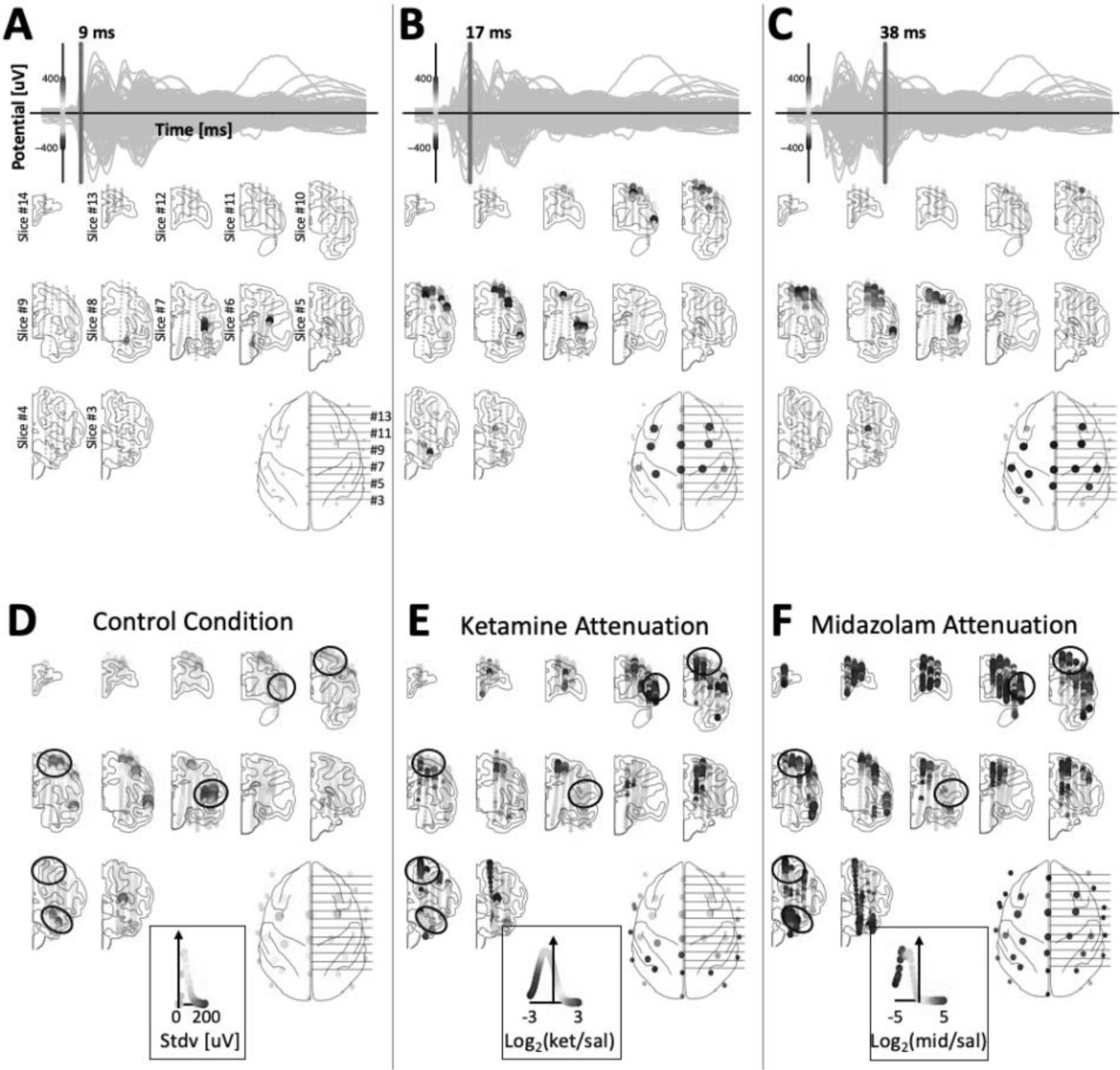
Spatial distribution of auditory evoked electric fields. (**A**–**C**) Hemisphere-wide auditory evoked electric fields. The top of each panel features a butterfly plot with time-resolved activity of all channels; the red line indicates the time-point displayed in each panel. The mapping of color to potential is indicated on the y-axis of the butterfly plot. The first activation of auditory cortex occurs ∼9 ms after sound onset (slice #7). Activity in motor and pre-frontal cortex reaches its first surface-positive peak 17 ms after sound onset (slices #8&9). A second peak with inverted polarity emerges around 38 ms after sound onset. Note the dipole-like structure of the fields in auditory and motor cortex. For visualization purposes, the evoked potentials at the EEG electrodes were multiplied by a factor of 2. (**D**) A map of the strength of the evoked fields measured as the standard deviation of the time-resolved evoked responses over the first 250 ms of the response. The inset depicts the distribution of evoked field strength across the hemisphere and defines the relationship between color and field strength (0uV: gray, 200uV: red). (**E**) The attenuation of evoked responses after administration of a subanesthetic dose of Ketamine. Note that responses in auditory cortex are not attenuated. (**F**) Attenuation of responses after a subanesthetic dose of midazolam is much more pronounced (note different color scale) and strongly attenuates responses in auditory cortex.

Consistent with earlier work ^22,47,54,55^, we identified dipole-like electric fields in and around auditory and motor cortex that are believed to contribute to auditory evoked EEG responses at the scalp, validating our system (**Fig. 2**A-C). Our results go beyond earlier work by simultaneously capturing electric fields across the entire hemisphere, rather than one or two-dimensional partial slices. It is noteworthy that the auditory evoked electric fields could be recorded from electrodes in adjacent white matter, or cortical and subcortical regions. This observation provides important insights into the nature of the LFP signals recorded with the MePhys system and show that the electrodes are not only sensitive to activity generated in the immediate proximity of the electrode, but also record activity from distal generators that likely reflect synchronized post-synaptic potentials across large populations of pyramidal neurons.

Note that the spatial maps in **Figure 2A-C** were computed from a single 5-minute-long recording session. This speaks to the richness and the high signal-to-noise ratio of the data. We have recorded more than 200 daily recording sessions, each lasting 2-3 hours and featuring several active and passive paradigms. Hence, the data and analyses highlighted here represent only a tiny fraction of the available data and focus more on the breadth of possible analyses, rather than an in-depth analysis of one specific topic.

#### Impact of consciousness-altering drugs on stimulus-evoked activity

We next studied how the meso- and macroscopic auditory evoked responses are impacted by a subanesthetic dose of ketamine that is believed to mimic certain aspects of psychosis. Our results confirmed earlier work that ketamine broadly attenuates auditory evoked EEG responses ^56–58^. By simultaneously recording auditory evoked fields across the entire hemisphere, we were now able to pinpoint these reductions to prefrontal, posterior parietal and motor cortex, as well as cerebellum (**Fig. 2E**). Interestingly, the effects of ketamine on auditory cortex itself were minimal, with no reduction in overall response amplitude, and only subtle changes in latency of peaks and troughs. This suggest that ketamine either attenuated the propagation of auditory information from auditory cortex to downstream regions, or selectively attenuated the responsiveness of all regions other than auditory cortex.

We then compared the effects of ketamine to those of midazolam, which also attenuates auditory evoked EEG responses ^57^ but does not mimic aspects of psychosis. In contrast to ketamine, midazolam led to strong attenuation of responses in auditory cortex. In addition, even stronger reductions of auditory evoked responses were observed across the entire hemisphere–responses in other cortical regions and cerebellum were almost eliminated (**Fig. 2F**).

### 2.3 Motor Output – Movement-related fields and beta-band desynchronization

To probe the ability of MePhys to measure hemisphere-wide motor signals, we used an auditory change detection task during which the animal released a lever when it detected a deviant target tone. Strong movement-related 18 Hz β-band desynchronization is visualized for two example regions, cerebellum (**CB**) and motor cortex (**F1**) (**Fig. 3A)**. **Figure 3B** shows the time-course of movement-related beta desynchronization for 7 brain regions in the auditory (**A1**, **R**), motor (**F1**, **F5**, **Cd**), and somatosensory system (**S1**). The strongest and earliest β-band desynchronization was observed in motor cortex, followed 25ms later by cerebellum and supplementary motor cortex (**Fig. 3B**). **Figures 3C&D** compare the timing and spatial distribution of the β-band desynchronization with that of the response-related evoked potentials. Though we observed a close spatial alignment between these two effects, β-band desynchronization emerged later, and lasted substantially longer than the motor-evoked responses. A main exception was region 7a, which showed strong movement-related responses but very little β-band desynchronization, highlighting the importance of simultaneous recording of activity from many different regions for full comprehension of neural dynamics underlying motor response.

**Figure 3.**
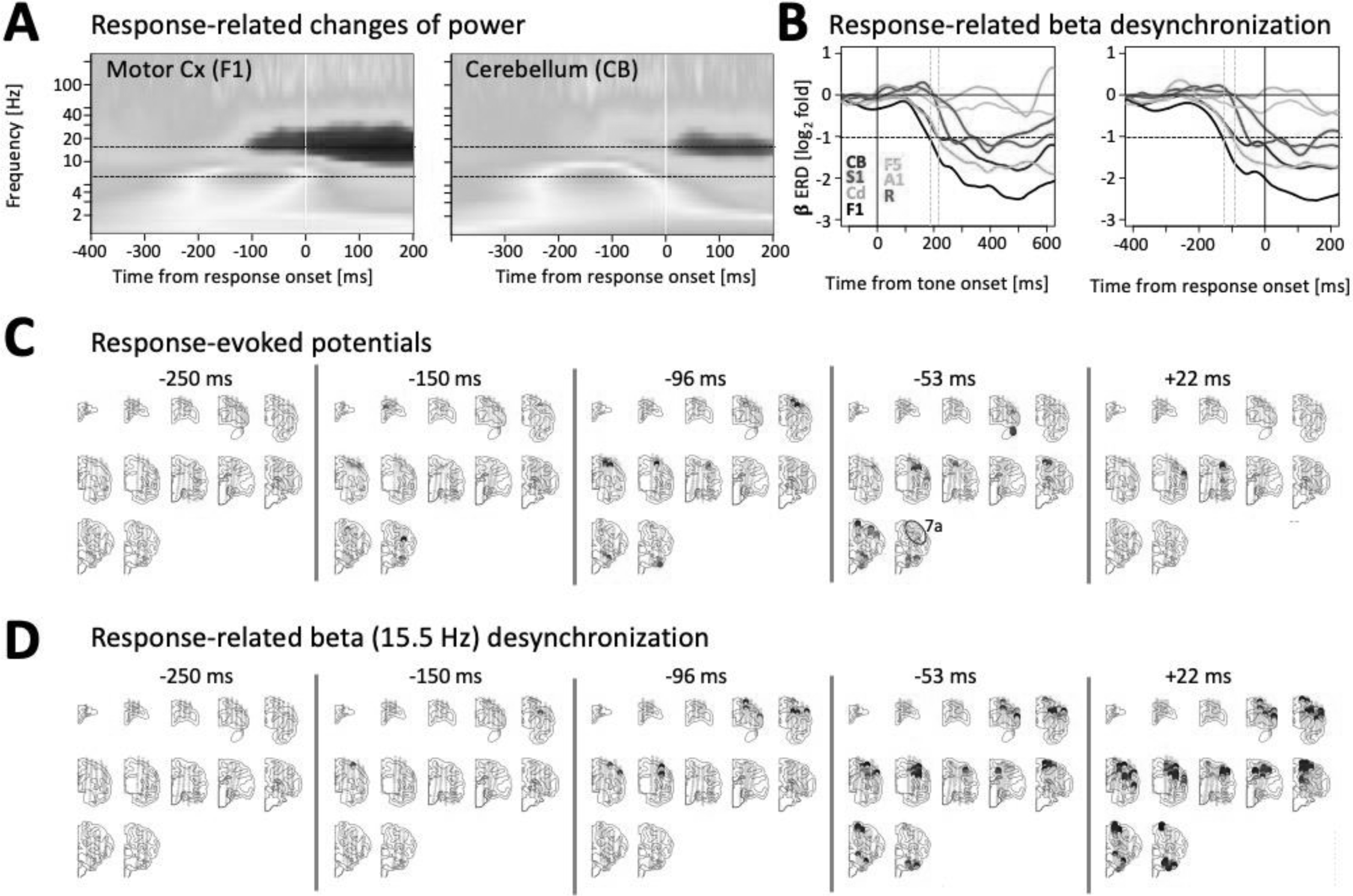
Movement-related fields and β-band desynchronization. (**A**) Wavelet decomposition of F1 and CB activity around the time of response onset shows an enhancement of alpha power and a subsequent reduction of beta power. (**B**) Timing of β-band response related desynchronization in 7 example regions (CB-Cerebellum; S1-Primary somatosensory Cx; Cd-Caudate; F1-primary motor Cx; F5-ventral premotor Cx; A1-primary auditory Cx; R-Rostral auditory Cx). The earliest drop in β power (defined as a reduction to less than half the power of control trials without a response) is observed in F1, ∼190 ms after tone onset/ 120 ms prior to response onset. Cerebellum and F5 had latencies that were ∼25 ms longer than F1. Somatosensory cortex reached criterion ∼50 ms prior to response onset. (**C**&**D**) Spatial maps of response-related evoked potentials and response-related β-band desynchronization at 5 different times relative to response onset.

### 2.4 Intrinsic Activity

To highlight how MePhys can help understand intrinsic activity, we transitioned to resting state recordings during 3-5-minute-long blocks without presentation of stimuli or tasks, both at baseline and after injection of sedative doses of (i) ketamine and (ii) midazolam. A frequency analysis during rest revealed spectra with a wide range of different shapes that varied systematically with electrode location (**Fig. 4** & **Suppl. Fig. 5**, black lines). Many spectra were dominated by a single peak. In other cases, we observed two or three distinct peaks, while others followed a 1/f – type decay. For each channel we defined the dominant frequency as the frequency with the highest power. During the control condition, the dominant frequency in occipital and parietal cortex was centered in the delta band around 3 Hz (**Fig. 4B**, left panel). The dominant frequency in frontal cortex, basal ganglia and cerebellum was centered in the low beta band around 12 Hz. Finally, there were numerous contacts in prefrontal, orbitofrontal and anterior temporal cortex with a dominant frequency in the high theta (or low alpha) range around 8 Hz. We did not identify any contacts with dominant frequencies above 25 Hz. However, contacts in and around ventral striatum, amygdala, and piriform cortex showed a narrow and distinct secondary peak in the gamma range centered on 40 Hz (**Fig. 4B viii**).

**Figure 4.**
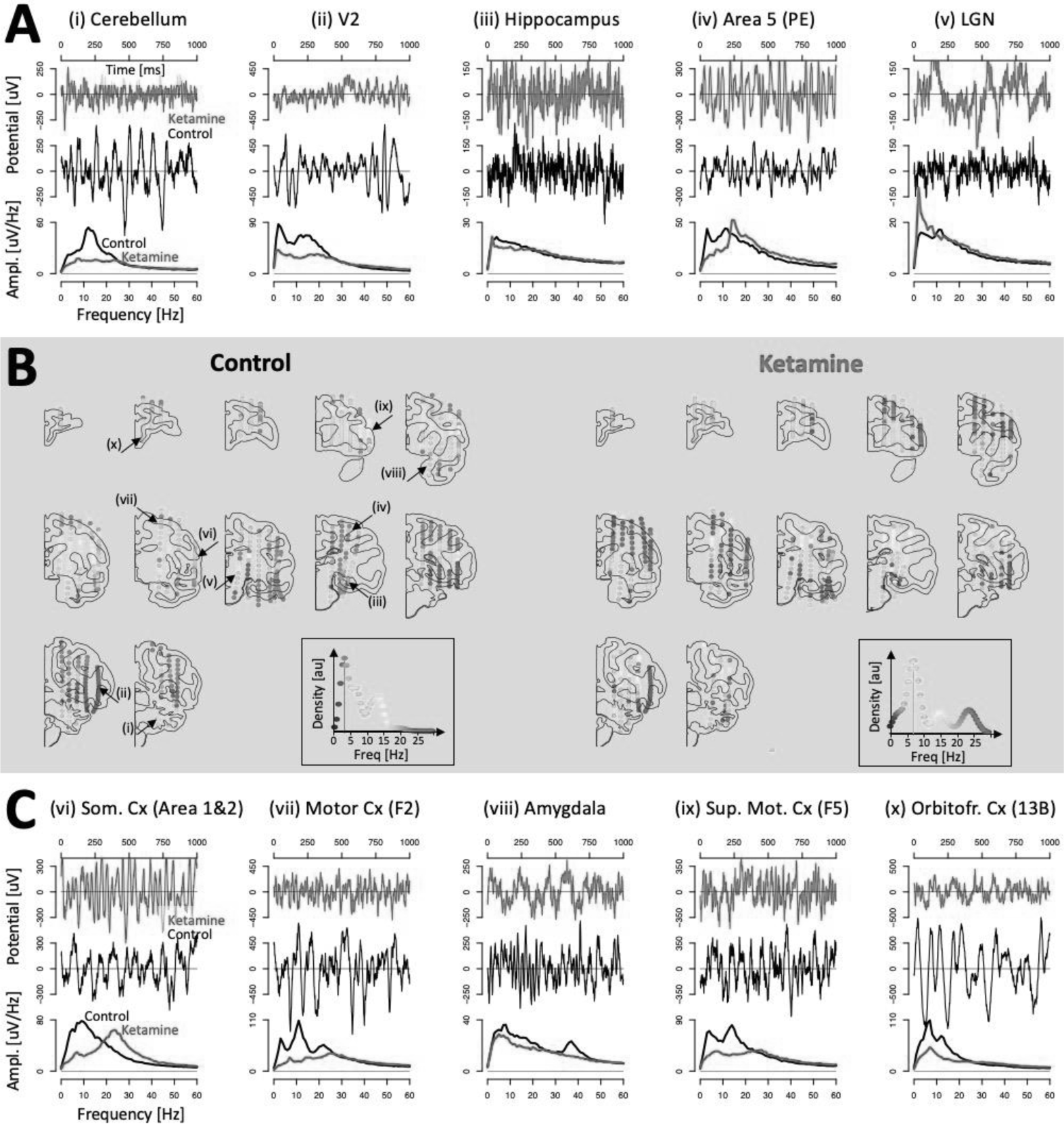
Distribution of dominant frequencies in the monkey brain. (**A**&**C**) Example traces and amplitude spectra for 10 channels during control (black) and after injection of ketamine (red). (**B**) Distribution of dominant frequency across the hemisphere during control (left) and after ketamine (right). Dominant frequency is color coded (Inset, frequency on the x-axis matches to color of the dots: blue [∼3Hz] – green [∼10 Hz] – yellow [∼15 Hz] – red [∼25 Hz]). The y-axis of the inset indicates the relative density of channels with the corresponding dominant frequency. During control, the mode of the distribution of dominant frequencies is at ∼3.5 Hz (blue vertical line). This peak is carried mostly by channels in occipital and parietal cortex. Channels in motor cortex and cerebellum are dominated by low beta. Channels in and around orbitofrontal cortex are dominated by low alpha. Overall, ketamine reduced power in lower frequency bands and the distribution of dominant frequencies shifted upwards (cyan vertical bar at ∼6.5 Hz). In rare instances, we see the emergence of new peaks (e.g., somato-sensory cortex or LGN).

The key effect of ketamine was to reduce power, especially for frequencies below ∼20 Hz; ketamine also flattened previously identified spectral peaks (**Fig. 4** & **Suppl. Fig. 5**, red lines). These effects were particularly striking in the cerebellum and motor cortex. Ketamine also eliminated the 40 Hz peak in and around the amygdala (**Fig.4C viii**). In contrast, the most striking effect of midazolam was the emergence and/or enhancement of ∼20 Hz peaks across large swaths of prefrontal cortex (**Suppl. Fig. 6B**, right panel, and **Suppl. Fig. 6C** ix & x). Midazolam also tended to flatten out peaks of the spectra, but the effects were less pronounced than for ketamine (**Suppl. Fig. 6**). For example, the 40 Hz oscillations in the amygdala that were abolished by ketamine were not affected by midazolam.

### 2.5 Intrinsic Coupling

One of the key applications of MePhys is description of large-scale hemisphere-wide functional networks with high spectro-temporal resolution that is not available in classical fMRI-based functional connectivity analyses. **Figure 5A** visualizes four such functional connectivity maps using example seed channels in the cerebellum, auditory cortex, amygdala, and the temporo-parietal junction.

**Figure 5.**
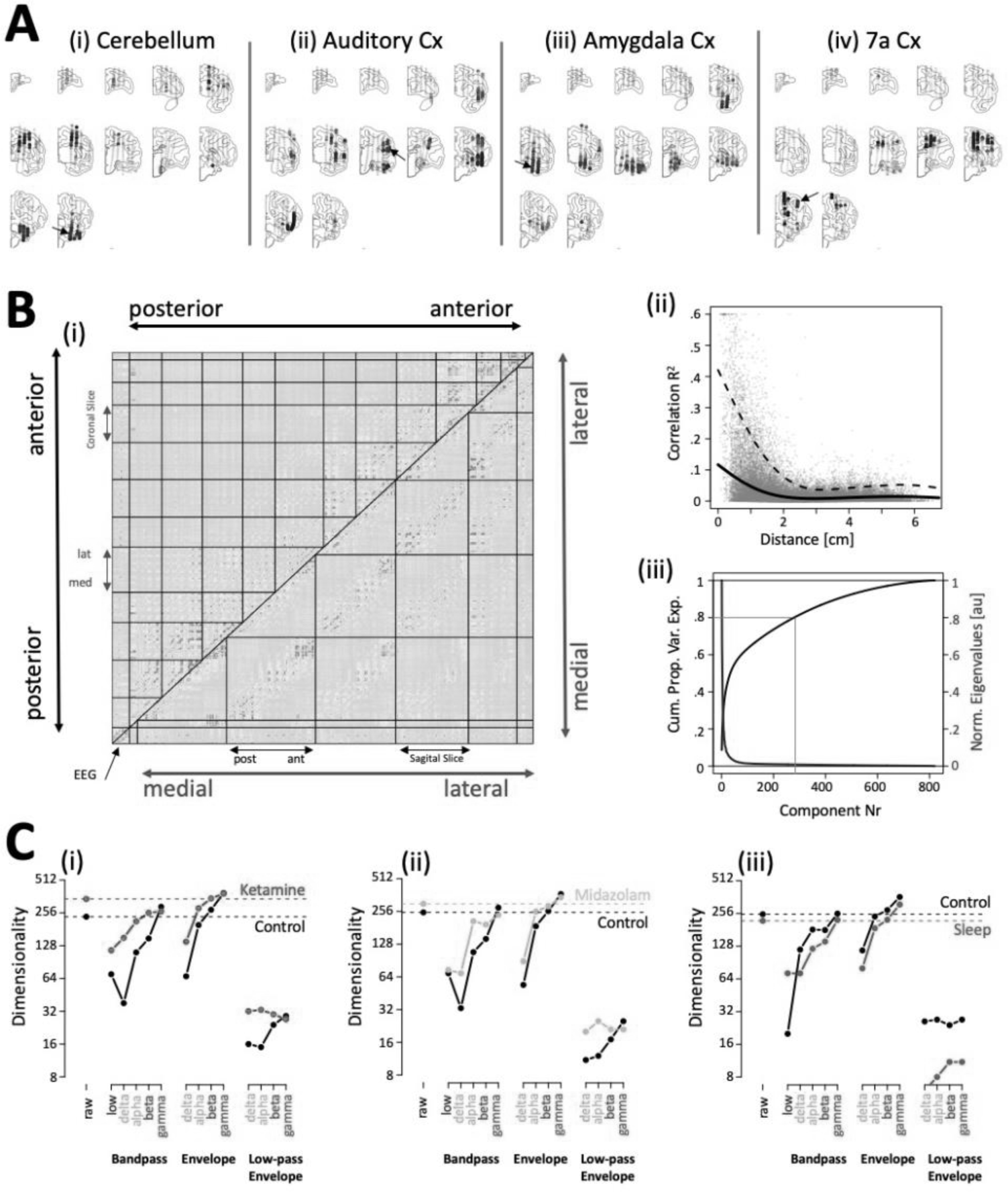
Intrinsic coupling. (**A**) Each panel shows the z-scored temporal correlation of one seed channel (black arrow) with all other channels. (**i**) The cerebellum seed confirms the existence of a functional link between cerebellum and motor cortex 59. The rapid transition between channels with strong positive and negative correlations in motor cortex reflects their location either above or below a dipole in motor cortex. (**ii**) The seed in auditory cortex identified functional connections along the entire superior temporal plane, extending all the way to the temporal pole as well as with motor cortex and ventrolateral prefrontal cortex. (**iii**) The amygdala seed revealed the expected functional connections to hippocampal areas and the ventral visual path, as well as the ventral striatum. (**iv**) The seed in area 7a revealed more complicated map with functional connections in hippocampal regions, dorsal prefrontal cortex and anterior Insula. (**B**) (**i**) Matrix of temporal correlations of the raw data for all pairs of channels. Intracranial channels are grouped either by coronal or sagittal slice (above and below the main diagonal, respectively). Within each slice, channels are sorted from medial to lateral or posterior to anterior, respectively. Bands of positive correlation along the main and secondary diagonals indicate stronger correlation for channels that are close to each other. (**ii**) Correlation plotted as a function of distance between each pair of electrodes. (**iii**) Normalized eigenvalues and cumulative percent variance explained by successive principal components. Dimensionality was defined as the number of components needed to explain 80% of the variance (gray lines) **(C)** Dimensionality (y-axis) of functional connectomes separated by carrier frequencies and mode before and after injection of ketamine (left) midazolam (middle) and falling asleep (right).

As a next step, we computed the full matrix of temporal correlations for all >300,000 simultaneously recorded pairs of channels. Following the nomenclature in the fMRI field ^7^, we refer to this matrix as the (mesoscopic) functional connectome. **Figure 5B** visualizes the functional connectome for the raw data in the control condition prior to injection of ketamine. The analysis identified a wide range of positive and negative correlations with meaningful spatial variation. The average value of the correlations (spearman correlation *r*^2^) decreased with distance, dropping close to zero at ∼2 cm (**Fig. 5Bii**). We defined the dimensionality of the connectome as the number of components needed to explain 80% of the variance (**Fig. 5Biii**).

Because brain function and communication has been suggested to occur in specific frequency bands, we filtered raw LFP data in five frequency bands: low (0.1 – 1 Hz), delta (2 – 4 Hz), theta/low alpha (5 – 10 Hz), low beta/mu (10 – 20 Hz) and low gamma (20 – 40 Hz). Frequencies below 0.1 Hz did not contain any meaningful signal because of the hardware filter of the amplifier. We also calculated the envelopes for four band-passed data sets (delta, alpha, beta, gamma) by low-pass filtering the full-wave rectified band-passed data. It has been suggested that power fluctuations in certain frequency bands may be a homolog of BOLD fluctuations ^5^. Hence, we created two versions of the envelopes with either a high- or lowpass filter of 0.1 Hz.

Using a seed-channel in cerebellum as an example, we show that the functional connectivity maps indeed depend on the 14 different filter settings (**Suppl. Fig. 7**). We furthermore leveraged the high temporal resolution of MePhys to quantify the temporal lag between different brain regions. For example, our results show that beta envelope in motor cortex leads cerebellum by ∼2 ms (**Suppl. Fig. 8**). Such spectral and temporal specificity cannot be identified using methods that lack millisecond temporal resolution.

We next computed the functional connectomes for the band-passed data sets and their envelopes. Dimensionality of the connectomes varied systematically as a function of carrier frequency, mode, and experimental condition (**Fig. 5C**). (1) With few exceptions, dimensionality increased with carrier frequency. (2) Dimensionality of the low-passed envelopes is markedly lower than the bands and envelopes. (3) Ketamine increased dimensionality for almost all carrier frequencies and modes. A similar but smaller amplitude effect was observed for midazolam. In contrast to ketamine and midazolam, sleep decreased the dimensionality of the connectomes.

### 2.6 Intrinsic States

Another promising application for MePhys is the ability to study if and how the brain transitions through different intrinsic states over time. The low dimensionality of the low-passed envelopes may reflect a small number of intrinsic states mediated by slow-acting transmitter systems (neuropeptides), or slowly fluctuating physiological (hunger, thirst, etc), emotional (fear, anxiety, etc) or cognitive states (attention, arousal, etc). We concatenated the four low-passed envelopes and calculated spatial correlations between all pairs of time-points. As the temporal distance between two sets of maps increases, their similarity tends to decrease. However, our results also show that between periods of change, the brain settles into relatively stable states for periods on the order of tens of seconds (**Fig. 6A&B**). The patches of strong positive correlations visible off of the main diagonal suggests that the same stable patterns tend to recur at different points in time. The block-like structure also suggests rapid transitions between different intrinsic states. Interestingly, this entire structure is almost completely abolished following the injection of ketamine (**Fig 6C**). In contrast, the injection of midazolam had no effect on the emergence and recurrence of stable states (**Fig 6D**).

**Figure 6.**
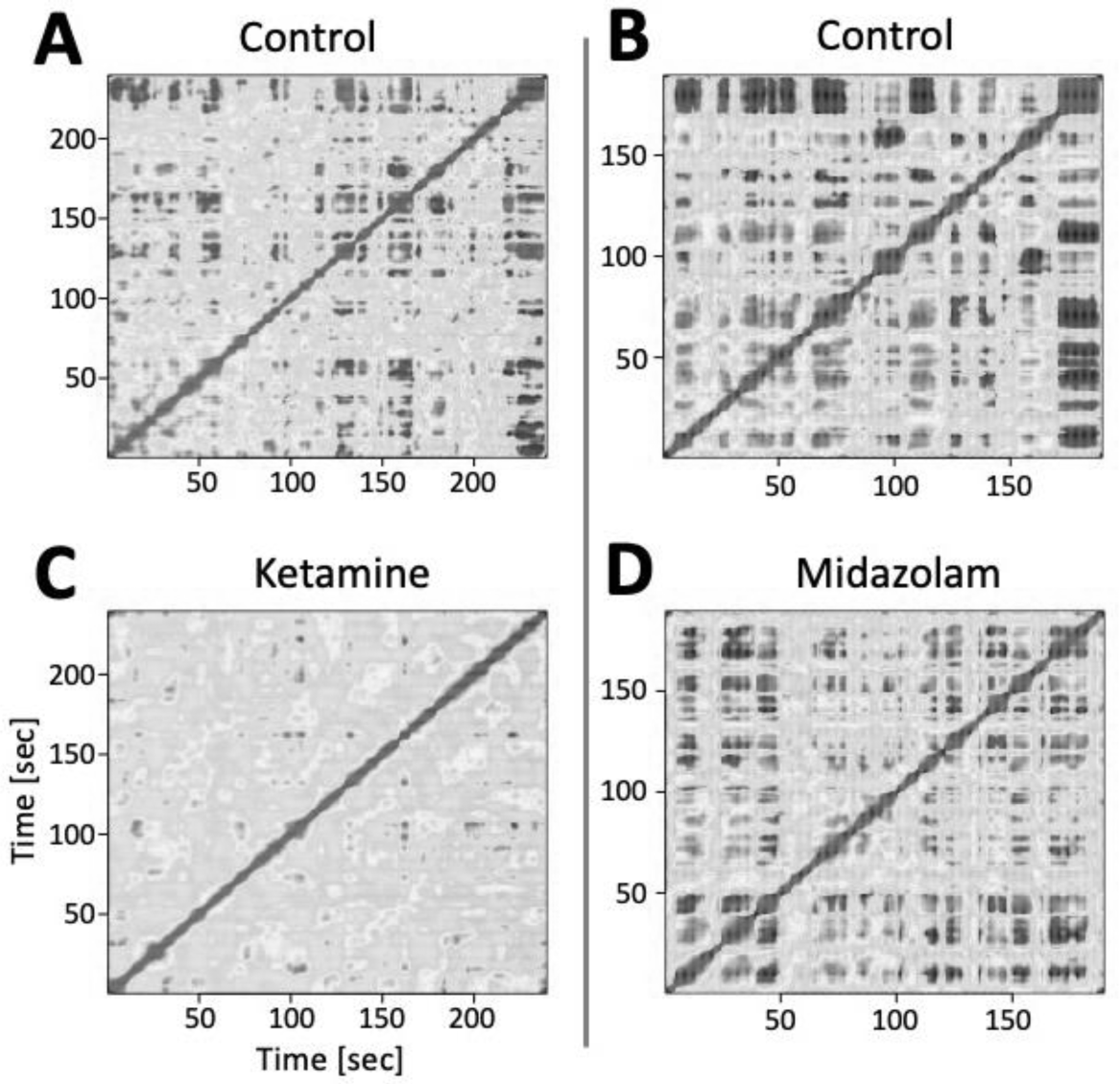
Ketamine disrupts the emergence of stable recurrent brain-wide states. Spatial correlation matrices during rest computed from all four low-pass envelopes simultaneously before and after ketamine (**A**&**C**) before and after midazolam (**B**&**D**). Color indicates the similarity of the spatial maps at different time points in the recording session. The thickening of the main diagonal into square-like patches along the main diagonal indicate that between periods of change, the brain settles into relatively stable periods that can last tens of seconds. The patches with strong positive correlations off of the main diagonal indicate that the same stable states occur at different points in time. Their square-like shape indicates relatively brisk transitions between states. The emergence of the stable states is largely abolished by ketamine (**C**), but not midazolam (**D**).

## 3. Discussion

We have developed the first volumetric mesoscopic electrophysiology platform by chronically implanting an approximately regular three-dimensional grid of electrode contacts across an entire monkey hemisphere, including cerebrum, cerebellum, diencephalon, and parts of the mesencephalon. By sampling LFPs in a three-dimensional grid, we created a new imaging modality, mesoscopic electrophysiology, or MePhys, that combines the millisecond temporal resolution of electrophysiology with the large field of view and millimeter spatial resolution of fMRI.

### Hemisphere-wide spatial maps

A key strength is that MePhys yields hemisphere-wide spatial maps of any LFP-based feature, such as sound-evoked electric fields (**Fig. 2**), response-related beta-band desynchronization (**Fig. 3B&C**) or resting-state spectra (**Fig. 4B**, **Suppl. Fig 7B**). Such hemisphere-wide maps provide important spatial context that is lacking when recording from only one or few brain regions. The feature maps varied gradually and systematically over space and followed anatomical landmarks such as the border between cerebellum and occipital cortex (**Fig. 2–5**).

The spatial smoothness is likely caused by two factors: (i) functional similarity of computations in close-by patches of cortex, and (ii) the low-pass properties of neural tissue ^20^ that allow LFPs to propagate and average across space. Together, this can enable the emergence of so-called generators, i.e., electric fields created by synchronized post-synaptic currents in large populations of pyramidal cells. Due to the anisotropic anatomy of pyramidal cells, these generators can give rise to a dipole-like spatial arrangement of current sources and sinks (**Fig. 2A-C**), that can propagate all the way to the scalp where they can be recorded as EEG potentials ^22–27^.

The signals recorded on each MePhys electrode likely reflect a mix of many different EEG generators. **Figure 2C**, for example, identifies at least two dipole-like electric fields, one in auditory, one in motor cortex. Depending on their distance from these two generators, each electrode will reflect activity of these generators with different degree and polarity. Changes in the location of an electrode will change the exact contribution of each generator. For example, moving an electrode from below to above the superior temporal plane will change the polarity of the contribution of the generator in auditory cortex. However, it will still reflect the same two generators active at that time. This is critical, because it suggests that – just like EEG results that are largely independent of the exact montage used– key findings from MePhys are independent of the exact location of the electrode contacts on the grid. Hence, while MePhys clearly does not record from every neuron, cortical layer or even from every region, it can certainly capture activity from EEG generators located anywhere in the hemisphere. Furthermore, it can provide much higher spatial resolution and signal-to-noise ratio compared with EEG or ECoG electrodes, located outside the skull or brain parenchyma, respectively. Thus, MePhys allows us to draw conclusions about the activity of generators in the entire hemisphere at the mesoscopic scale, a unique capability not available from other existing recording paradigms.

#### Intrinsic functional connectivity

A second key strength is that MePhys simultaneously records data from more than 300,000 pairs of electrodes distributed across the entire hemisphere. The large number of pairs and their wide field of view provides substantially more information than functional connectivity analyses using microscopic recordings that typically target no more than a handful of local regions. For example, our seed-based functional connectivity maps confirmed a very localized connection between cerebellum and motor cortex (**Fig. 5Ai**) that has previously been identified with fMRI ^59^. In addition, MePhys was able to add spectro-temporal information by firmly localizing the functional connection to the mu/low beta band (**Suppl. Fig. 7**), and by quantifying the temporal lag to be on the order of 2 milliseconds (**Suppl. Fig. 8**).

We were also able to quantify key properties of the hemisphere-wide functional connectome as a function of carrier frequency (delta, alpha, beta, gamma) and mode (band vs envelope vs low-pass envelope). We identified the strongest correlations and the lowest dimensionality for the low-passed envelopes for the delta and alpha carrier frequencies. These signals seem to emerge as the most likely homologs of the slow BOLD fluctuations that underly fMRI functional connectivity measures based on their frequency below 0.1 Hz, as well as their low dimensionality and signal strength. Being sensitive to both fast and slow time scales, MePhys thus has the potential to help us understand if and how large-scale networks of functional connectivity at fast time-scales are nested into and modified by the dynamics of intrinsic activity at slower timescales.

#### Hemisphere-wide intrinsic states

Using spatial correlations of the low-passed envelopes, MePhys was able to show the emergence and recurrence of stable large-scale states that can last on the order of tens of seconds (**Fig. 6**). Ongoing work is aimed at creating a library of LFP-derived intrinsic states and reliably linking them the activity of slow-acting neuropeptides ^60^ or to psychological constructs such as hunger, thirst, arousal, fear, motivation, or attention.

#### Open science

Centralized brain observatories for large-scale recordings in mice and non-human primates will likely play an increasingly important role in neuroscience ^61^. Their goal is to make mouse or non-human primate data more easily accessible for the computational neuroscience or human neuroimaging communities. To date, however, non-human primate brain observatories don’t exist, and non-human primate data sets remain extremely time-consuming and expensive to acquire. We believe that MePhys is an ideal mesoscopic imaging modality for the non-human primate brain observatories of the future. Daily collection of MePhys data requires minimal setup time and can be performed by a dedicated technician. Because electrodes are chronically implanted and spread out to record LFPs from across an entire hemisphere, they don’t need to be moved between sessions to sample from different neurons and/or different parts of the brain. As a result, MePhys can provide comprehensive data sets for a given task within days or weeks, rather than years. The main limiting factor that determines how quickly animals can transition between paradigms is thus the behavioral training of the animals. The mesoscopic results can then provide a strong rationale for in-depth single-cell recordings in brain regions with interesting LFP responses.

#### Recording during learning

The ability to record comprehensive MePhys datasets quickly also paves the way to track changes of brain function across different stages of learning. This adds a critical new angle to monkey electrophysiology experiments which are typically performed over the course of years in highly overtrained animals.

#### Translational relevance

The non-human primate is a crucial element of a translational pipeline designed to bring novel therapeutic approaches from the bench to the bedside. The large disconnect between single-cell recordings in monkeys and EEG/MEG or fMRI recordings in humans is a weak point of this pipeline. By combining the best features of both EEG and fMRI into a single method in the non-human primate, MePhys will be able to facilitate the comparison between macroscopic findings in humans and microscopic single-cell recordings non-human primates, and thus the translation of findings from rodents, via monkeys to humans, and vice versa.

#### Limitations

The strengths of MePhys of course come with their own specific limitations. (**i**) Over the long run, the chronic nature of the implant is not conducive to single cell recordings. While we observed stable single-cell activity with high signal-to-noise ratio over the first two months, most of them have since transitioned to multi-unit activity. (**ii**) In contrast to acute or semi-chronic recording systems ^3,17^ (Gray Matter SC drives), the position of MePhys electrodes can’t be adjusted. (**iii**) The spatial density of electrode contacts will always be below what can be achieved when targeting all electrodes to a smaller portion of the brain. Hence, mesoscopic electrophysiology does not compete with classic microscopic electrophysiology. Instead, MePhys complements existing micro- and macroscopic methods and will create synergy by bridging some of the gaps between them. (**iv**) In contrast to fMRI or EEG, MePhys is an invasive methodology that causes damage during implantation. The T-Probes used for MePhys are thinner, but otherwise very similar to SEEG electrodes that are used for epilepsy monitoring in the clinic. Interestingly, neurosurgeons are routinely implanting tens of SEEG electrodes, including in pediatric patients, with few side effects. The only cautionary note is that the side-effects of SEEG electrodes, while rare, tend to be more severe (e.g., rupturing of a vessel) than side-effects of epidural ECoG grids (e.g., infections). This matches our experience with the MePhys surgeries. During the last surgery, we implanted 30 shafts and observed no post-surgical behavioral impact whatsoever. At this point, it remains to be determined if the damage incurred by chronic, i.e., one-time, implantation of many MePhys shafts is more problematic than the repeated acute implantation of a few shafts for classical microscopic electrophysiology.

#### Opportunity

While our first prototype provides a new window into non-human primate brain function, MePhys has not reached its full potential. (**i**) A re-design of the backends of the probes has already made them more robust, thus removing the main cause of signal loss in our prototype. (**ii**) A redesign of the MePhys platform will provide better access to occipital cortex and midline structures that were under-sampled in our prototype. (**iii**) Improved design and manufacturing of the probes will prevent the unnecessary tissue damage that was caused by bending and/or bucking probes in the first implant surgeries. (**iv**) Brain damage can be further minimized once it is possible to replace stainless steel with flexible electrode shafts that move with the brain. (**v**) Most importantly, channel count and electrode density of the implants will continue to increase in line with ongoing efforts to miniaturize electrodes and recording equipment. The next generation MePhys systems is currently designed to feature between 120 and 155 electrode shafts for a total of up to 3200 intracranial contacts. We expect that within 5 years we will be able to increase the number of electrodes along each shaft to 256, the density of electrode shafts in the antero-posterior direction from 5 to 3 mm, and from 4 to 3 mm in the medio-lateral direction all while maintaining an entire hemisphere as a field of view. Our mid-term goal is to be able to implant 1024 channel probes such as Neuropixels ^16^ or SiNAPS probes ^18^ (Plexon or Neuronexus) at each location. (**vi**) As electrode density along each shaft increases, it will eventually be possible to compute current-source densities with laminar resolution across the entire brain, thus pushing the spatial resolution of MePhys closer to the microscopic scale. (**vii**) In addition to the technical improvements discussed above, future work will apply more sophisticated analytical approaches.

In summary, we believe that mesoscopic electrophysiology is a unique and versatile addition to the rapidly expanding toolkit of non-human primate electrophysiology. Key hallmarks of MePhys will be (**i**) the spatial scope on the entire hemisphere or brain, (**ii**) a temporal resolution on the order of micro-seconds, (**iii**) a relatively high spatial resolution that increases in line with technological advances to miniaturize components, and (**iv**) the ability to record comprehensive data sets quickly at different stages of learning, not just in overtrained animals.

## 4. Methods

### 4.1 Subjects

The experiments were performed on 1 adult male macaque monkeys (macaca mulatta). The treatment of the monkey was in accordance with the guidelines set by the U.S. Department of Health and Human Services (National Institutes of Health) for the care and use of laboratory animals. All methods were approved by the Institutional Animal Care and Use Committee at the University of Pittsburgh.

### 4.2 MePhys Platform

The MePhys platform consists of three rugged structural elements (*‘crown’*, *‘wall’*, and *‘cap’*), two internal elements that facilitate the implantation of the macroscopic and mesoscopic electrodes (*‘spider’* and *‘grid’*), as well as the head-stage (Spikegadget) and its *‘baseplate’* (**Fig. 1**). *Crown* and *wall* were initially designed in house using solidworks, and then refined and manufactured by Graymatter from PEEK using a 5-degree-of-freedom CNC machine. The *crown* is the main structural element that is anchored to the scull with metabond and a ring of ceramic screws around the outer margin. It forms a tight seal between the soft tissue on the outside, and the inside of the implant which consists of bone covered in metabond and acrylic. The *wall* is the second structural element that increases the height of the implant and provides the *‘wings’* for head-fixation.

The role of the *spider* and the *grid* is to determine the location of the EEG and intracranial electrode shafts, respectively. Both parts were designed in house using Fusion360 and manufactured from SOMOS resin on a VIPER^TM^ SLA printer. The top of the *spider* is the mounting point for the 32-channel omnetics connector of the EEG electrodes. The arms of the *spider* guide the wires from the connector to the electrodes without getting in the way of the to-be-implanted depth electrodes. The holes in the *grid* are the mounting points for the guide pins and thus determined the locations of the mesoscopic electrode shafts. The holes were spaced out by 5 and 4 mm in the antero-posterior and medio-lateral direction, respectively. The inner diameter of each grid-hole was 1.5 mm. The *baseplate* provided a stable platform for the Spikegadets headstage, and was designed to align the mini-HDMI connector with the corresponding opening in the cap. This allowed us to record from all channels without taking off the cap. The cap was 3D printed with Nylon by Graymatter. In the latest iteration, the cap is split into two parts to provide easy and flexible access to the amplifier and the connectors without having to take off the entire cap.

### 4.3 MePhys Guide Pins

The guide pins are one of the most important functional elements of the MePhys system. Their role is (a) to allow all 62 electrode shafts to penetrate the dura without buckling or breaking and (b) to subsequently keep infections from entering the brain along any of the electrode shafts. The guide pins were designed in house in Fusion360 and manufactured from SOMOS resin on a VIPER^TM^ SLA printer. The outer diameter of the guide-pins was 1.45 mm allowing them to slide into the 1.5 mm holes in the *grid*. To help the electrodes penetrate the dura without buckling, the guide pins featured a very thin inner diameter (500 µm), that narrowed even further at the tip (335 µm). While the diameter of the stainless steel tubes of the electrode shafts was 260 µm, the actual diameter of the finished electrodes was a bit larger. Similarly, the actual inner diameter of the printed guide pins was smaller than the nominal 335 µm. As a result, the bottom of the guide pins provided extremely close lateral support and prevented any lateral movement during penetration. Furthermore, the length of the pins was custom designed for each grid hole to lightly touch and dimple the underlying dura, thus facilitating the penetration (**Suppl. Fig 2**). This was especially important for the most lateral shafts that had to penetrate the dura at an oblique angle. In addition, we also used puncture pins with identical dimensions to pre-puncture the dura in the exact location that the electrodes would later be inserted. Using this approach, all electrode shafts penetrated the dura without problems.

Several steps were taken to prevent infections from entering the brain along the electrode shafts or the outside of the guide pins. First, the guide pins were sterilized in a hydrogen peroxide sterilizer. Using fully sterile procedures, the sterilized pins were backfilled with sterile grease, and the tip was closed with a drop of KwikSil. Both approaches have been used successfully in the Graymatter SC system. Prior to being implanted, the prepared guide pins were again sterilized using hydrogen peroxide. To prevent infections from entering along the outside of the guide pins, they were cemented into the grid-hole during the surgery using metabond. The guide pins also featured an array of 4 small holes located 5 mm from the tip that allowed sterile grease to be pushed out of the guide pin when the electrode was inserted (**Suppl. Fig. 2**E). This created a ring of sterile grease in the tight space between the inside of the grid hole and the outside of the guide pin.

### 4.4 MePhys platform surgery

The MePhys system was implanted in a series of 5 surgeries distributed across a period of 8 months. All surgeries were performed under the highest standards of sterility, given that it would be impossible to treat any infection without removing the entire implant. Goal of the first surgery was to implant the MePhys platform and the EEG electrodes. Prior to the surgery, we prepared the EEG electrodes as well as the omnetics connector and fastened them to the spider. Both spider and EEG connector were mounted into the crown and sterilized. During the surgery we exposed the skull, removed any residual tissue, and sealed the pores in the bone with copalite varnish. *Crown* and *spider* were positioned on the skull using a stereotactic arm and cemented in place with Metabond. Next, the holes for the ceramic screws were drilled and taped. The ceramic screws were then fastened into the holes around the outside diameter of the *crown*. A second layer of Metabond was applied from inside and outside of the *crown* to create a tight seal. We then drilled the holes for the EEG electrodes into the skull and cemented them in place with small drops of Metabond. Additional ceramic screws were then added in free spaces over the left hemisphere to increase the connection between skull and implant. Next, we installed the *wall* and fixed it in place with set-screws. The gaps between the *crown* and *wall* were filled with a thin layer of Dowsil that had been applied prior to combining the two elements. The *grid* was positioned over the right hemisphere and cemented in place by filling the space underneath it with dental acrylic. The ceramic screws over the left hemisphere were covered with additional layers of dental cement. After inserting the *baseplate* over the left hemisphere, the implant was covered with a flat cap before recovering the animal.

### 4.5 Electrode implantation

After the initial surgery, a CT image was collected to determine the actual location of the platform and the exact locations of the grid holes. Based on this image we determined that 58 of the 72 grid holes were suitable for electrode implantation. Eleven of the grid holes were excluded because they were closer to the anterior cerebral artery in the longitudinal fissure than intended. The two grid holes in the most posterior slice were excluded because drilling through them might have damaged some of the EEG wires that seemed to have moved around during the surgery. One grid hole was excluded because it was near a fault in the underlying dental acrylic, which raised concerns about sterility. The electrode shafts were implanted over the course of the next four surgeries. Two shafts were implanted during the first electrode implant surgery. As we improved our implantation technique over time, we implanted 4, 21 and 31 electrode shafts during the next three surgeries. **Supplementary** Figure 1 shows pictures of the implant after the first (**Suppl. Fig. 1A-D**), third (**Suppl. Fig. 1E**) and fourth surgery (**Suppl. Fig. 2F**).

Because the back-ends of already implanted electrode shafts cannot be sterilized in place, it was increasingly more challenging to create a sterile field for subsequent surgeries. Our solution was to unplug the connectors of all previously implanted electrodes and cover them under a sterile custom-designed cap. The approached succeeded at creating a sterile field, but we lost functionality of several electrodes in the process of repeatedly unplugging and re-plugging them before and after surgeries.

To implant an individual depth electrode into a grid-hole, we first drilled through the underlying acrylic and bone (**Suppl. Fig. 2A&B**). We then pre-punctured the dura using a puncture pin (**Suppl. Fig. 2C**) and cemented a custom-designed guide pin in place that lightly touched and dimpled the underlying dura (**Suppl. Fig. 2D**). Next, we inserted the electrode shaft into the guide pin and slowly pushed it through the dura and into the brain by hand until it reached its final position inside the guide pin (**Suppl. Fig. 2E**). Finally, the top of the guide-pins and the shaft of the electrodes were covered with 3D resin that was polymerized in place with a UV lamp during surgery (**Suppl. Fig. 2F**). Since the grid had been printed with the same resin, this provided an extremely tight and stable seal.

### 4.6 Yield and signal quality

Of the 57 electrode shafts that we implanted, 56 were fully functional after the respective surgery. However, 6 of them subsequently lost functionality due to mechanical problems with the electrode backend (**Suppl. Fig. 4**). In four cases, we can link the loss of signal to plugging and unplugging of the connector that was necessary for subsequent surgeries. In two instances, the exposed tops of the electrodes with the pig-tails ripped off. Other than these instances of mechanical failure, we have not noticed any degradation of the recorded local field potentials over time. As expected, we did lose isolated single cells over the first couple of months. However, we can still record stable multi-unit activity on many of the same channels more than four months after implantation.

### 4.7 Recording system

The MePhys system is built around the 1024 channel modular headstage from SpikeGadgets (SpikeGadgets LLC) ^62^. The headstage is small and light enough to be permanently enclosed in the MePhys platform on the left side of the animal’s head. It provides easy access to all 1024 channels via a single micro-HDMI connector at a sampling rate of 20 or 30 kHz. The SpikeGadets system also allows untethered recordings, a feature that we plan to take advantage of soon. The system consists of a maximum of 8 modular amplifier boards each supporting 128 channels via two 64-channel ZIF connectors. The T-Probes were connected to the headstage using flexible printed circuit board (PCB) cables that were designed in house and manufactured by PCBWay. The flex-PCB cables featured a 64 channel ZIF connector on one end that branched into 4 narrower cables, each with a female 18-channel omnetics connector that mated with the male omnetics connectors of the T-Probes. The exact layout of each of the 16 flex PCB cables was designed in Fusion360 to each match the location of a different set of 4 of the 62 planned electrode shafts. This custom design of the backend was critical to enable a coherent design of the whole system, including the implantation grid, the location of the headstages and the management of the flexible cables.

### 4.8 MePhys electrode layout

We designed our MePhys system to combine 32 macroscopic EEG electrodes with 992 intracranial electrode contacts distributed across 62 electrode shafts arranged on a regular grid in the horizontal plane over the right hemisphere. The grid was arranged into 14 coronal slices and featured between 2 and 7 grid holes per slice. Based on CT and MRI images we customized the electrode shafts for each grid hole to traverse the underlying Tel-, Met-, and Diencephalon, as well as parts of the Mesencephalon, but excluding the pons and medulla, and avoiding the Circle of Willis. The 16 electrode contacts on each shaft were spaced out regularly across the entire section of brain parenchyma traversed by the electrode shaft. The EEG electrodes were positioned in an approximately regular 2D array on the skull surface that was designed using our monkey 10-20 software package ^63^. EEG electrode locations on the right hemisphere were adjusted slightly so as not to interfere with the location of the intracranial electrode shafts.

### 4.9 Electrode locations

To confirm the intended locations of the electrode shafts and contacts, we acquired a post-surgical CT scan with all the implanted electrode shafts in place. In most cases, the electrode shafts were close to their intended locations (**Suppl. Fig. 3**). We observed a slight slant in some shafts which we believe was caused by minor pre-existing bends in the stainless-steel tubes. Two electrode shafts seemed to be entering the brain at an oblique angle, suggesting that the holes through the acrylic had not been drilled perfectly perpendicularly to the *grid*. This is likely the result of these holes being drilled by hand. There were also some subtle discrepancies in the vertical position of the probes. These vertical discrepancies arose from the fact that all electrode shafts were between ∼3 to 13 mm longer than specified, due to a miscommunication with the manufacturer. This discrepancy was only noticed after the third electrode implant surgery and caused the affected electrodes to penetrate deeper than intended. In four extreme cases, the additional length caused the electrode shafts to come into contact with the bone on ventral side of the skull, and buckle. However, in most instances the electrode contacts just ended up a bit deeper than originally intended, causing the topmost contacts to be below the dorsal cortex, rather than above it (**Suppl. Fig. 3B**, slice #2 through #7; ap -1.15 through +1.15). Once we became aware of the problem, we corrected for the additional length by adding a custom-designed stopper in house. This improved the match between intended and actual electrode location for the 31 shafts implanted in the last surgery. However, the stopper tended to overcompensate and left some of the topmost electrode contacts outside of the brain parenchyma (**Suppl. Fig. 3B**, slice #8 through #14; ap +1.7 through +4.7). Unfortunately, the mechanical link between stopper and electrode shaft was weaker than for the original reinforcement tube in some cases. In combination with the electrode shafts extending up to 13 mm higher above the grid than intended, this left electrode back-ends more vulnerable forces generated by rapid head-movements and caused two of them to break off. We have since mechanically stabilized the remaining shafts using Loctite Foam.

### 4.10 Cranial EEG recordings

Details of the cranial EEG recordings have been reported previously ^64,65^. Briefly, 33 EEG electrodes manufactured in-house from medical grade titanium were implanted in 1mm deep, non-penetrating holes in the cranium. The electrodes were connected to a 36-pin Omnetics connector mounted in the left posterior aspect of the MePhys platform. The 33 EEG electrodes formed an approximately regularly-spaced grid on the skull covering roughly the same anatomy covered by the international 10-20 system ^63^. The position of EEG electrodes on the right hemisphere were adjusted to fall in between the grid holes for the intracranial electrode shafts.

### 4.11 Intracranial T-Probe Electrodes

The intracranial probes (T-Probes) were custom designed and hand-made multi-contact probes (NeuronElektrod Kft, distributed by Plexon Inc.). The dimensions of the T-Probes were determined based on a CT scan acquired after the implantation of the MePhys platform. Based on this CT image, we determined individual electrode specifications for each of the 62 grid holes based on the underlying anatomy. Two main parameters determined the layout of each electrode. (1) the length of the *active* portion of the shank towards the tip of the probe ‘*a*’, and the *blank* or passive portion towards the top of the probe ‘*b*’ (**Fig 1B**). The length of the blank portion of the probe was designed to traverse the distance from the surface of the grid to the inner surface of the dorsal aspect of the skull. The length of the active portion of the probes was chosen to traverse the entire underlying brain tissue, including Tel-, Met-, and Diencephalon, as well as parts of the mesencephalon, but excluding the pons and medulla, and avoiding the Circle of Willis. The 16 electrodes were spaced uniformly across the active portion of the shank, thus determining the inter-electrode spacing. Depending on the length of the active portion, inter-electrode distance along a shaft ranged between 2.7 and 0.4 mm. The very top of each shank contained a 9 mm long reinforced section with a pig-tail connector that terminated in an 18 channel omnetics connector.

From a technical point of view, T-probes are a modification of the popular V-and S-Probe. The requirement of site spacing was to be equidistant across the entire active portion of the shank rather than the standard pre-defined 30µm, 50µm, or 75µm, etc. Since the active portion of the shank was different for each probe, the required inter-electrode spacing also differed for each probe. The key engineering challenge of the T-Probes was to accommodate the large inter-electrode spacing that required the opening in the stainless steel tube to be considerably larger than in the standard V- and S-Probe and still keeping suitable strength and flexibility of the shaft. This required the development of new manufacturing devices and protocols. Thus, it presented a significant challenge to manufacture the large number of probes in the limited amount of time dictated by the constraints of the project. (The exact dimensions of each probe are only available after implantation of the base, and any delay would thus shorten the overall lifetime of the implant.)

### 4.12 Experimental Setup

All experiments were performed in a small (4’ wide by 4’ deep by 8’ high) sound-attenuating and electrically insulated recording booth (Eckel Noise Control Technology). The animal was positioned and head-fixed in a custom-made primate chair (Scientific Design). Neural signals were recorded with a 1024-channel digital amplifier system (*Spikegadget) at a sampling rate of* 20 kHz.

Experimental control was handled by a windows PC running an in-house modified version of the Matlab software-package *monkeylogic*. Sound files were generated prior to the experiments and presented by a sub-routine of the Matlab package *Psychtoolbox*. The sound-files were presented using the right audio-channel of a high-definition stereo PCI sound card (M-192 from M-Audiophile) operating at a sampling rate of 96 kHz and 24 bit resolution. The analog audio-signal was then amplified by a 300 Watt amplifier (QSC GX3). The amplified electric signals were converted to sound waves using two single element 4 inch full-range driver (Tang Band W4-1879) located 8 inches in front of the animal to the left and right of the screen.

### 4.13 Recording paradigms

#### Resting state paradigm (resting)

The resting state paradigm consisted of ∼3-5 minute long blocks during which the animal was not required to perform any task, and during which no external stimuli were presented. During that time, eye-position and pupil diameter were monitored with an infrared eye-tracker. Resting state data sets analyzed here were recorded before and after injection of subanesthetic doses of ketamine (1.9 mg/kg) and midazolam (0.39 mg/kg). In addition, we analyzed two resting state data sets recorded right before and after the animal fell asleep. These two recordings were made in the evening (6.30 pm) and night-time (11.00 pm). All other recordings were made in the context of his normal work routine which typically extends from late morning (10 am) to early afternoon (1.30 pm).

#### Passive auditory paradigm (RFmap noise)

In the RFmap noise paradigm, animals passively listened to brief 25 ms long pulses of white noise with a 5ms linear rise/fall time. The white noise bursts were presented at 5 intensity levels (46, 56, 66, 76, 86 dB SPL). Stimulus onset intervals ranged between 300 and 400 ms. The RFmap noise data sets analyzed here were recorded before and after injection of subanesthetic doses of ketamine (4.5 mg/kg) and midazolam (0.48 mg/kg).

#### Delayed frequency discrimination task (change detection)

The delayed frequency discrimination paradigm has been described in detail before ^66^. Briefly, the task measured the animals’ ability to discriminate the tonal frequency of sequentially occurring pure tone pips. Each trial consisted of up to 13 tones (60 dB SPL, 205 ms duration, 5 ms rise and fall time). 80% of trials contained one target tone of different frequency relative to the preceding standard tones (target-present trials). In the remaining 20% of trials, the ‘target’ tone was identical to the standard (catch trials). Animals moved a lever off-center to initiate a trial. On catch trials, animals were rewarded for not releasing the lever during the entire trial. In the ‘target-present trials’, they were rewarded for releasing the lever between 100 and 800 ms after target onset. For the analyses presented here, we focused on the trials in which the animal released the lever, as well as the same number of matched control trials without a lever release.

### 4.14 Preprocessing of local field potentials

Time-continuous raw data was filtered in MATLAB using a 256 point non-causal digital low-pass FIR filter (*firws* function from EEG-lab toolbox, Blackman Window, high-frequency cutoff: 400 Hz). The filtered data was then cut into short epochs around external events such as sound onset. The epoched data was then downsampled from from 20 to 1 kHz and saved as MATLAB data files. In the case of the resting state recordings which contained no external events, we split the data into continuous 1-second-long chunks that could either be analyzed as 1-second long trials, or concatenated to recover the time-continuous traces.

### 4.15 Analysis of auditory evoked potentials

The main analyses were performed using in-house analysis software package implemented in R ^67^. After reading data from all channels, we computed the averaged auditory evoked potentials as a function of the 5 different sound pressure levels. Time-resolved auditory evoked potentials were visualized in multi-plots in which the relative position of each panel corresponded approximately to the relative position of the channels in each slice. For visualization purposes, the potentials at EEG electrodes were scaled by a factor of 2. In addition, we presented data in the form of heat plot panels that visualized activity of the intracranial electrode contacts distributed across 12 coronal slices as well as the activity of the EEG electrodes distributed over the surface of the skull.

### 4.16 Analysis of resting state data

After loading the epoched LFP data of all functional channels, the data was re-referenced to the common reference of all intracranial channels. LFP amplitude spectra were computed using fast Fourier transformation in each of the 1 second long epochs using a 25 ms long cosine window taper. The spectra were then averaged across all trials. From the averaged spectra we extracted the dominant frequency as the frequency with the highest amplitude. The dominant frequency was then color coded and displayed as heat-maps.

#### Extraction of bands and envelopes

The epoched data was concatenated into a ∼3-5 minute long time-continuous version. The time-continuous data was bandpass filtered to extract activity in 5 frequency bands: low (0.1 – 1 Hz, 1^st^ order Butterworth filter), delta (2 – 4 Hz, 3^rd^ order), theta/(low) alpha (5 – 10 Hz, 3^rd^ order), (low) beta (10 – 20 Hz, 3^rd^ order) and gamma (20 – 40 Hz, 3^rd^ order). Envelopes were computed by full-wave rectifying the band-passed signals and band-pass filtering the resulting time-series with a 1^st^ order Butterworth filter: delta envelope (0.1 – 1 Hz), theta/low alpha envelope (0.1 – 5 Hz), beta envelope (0.1 – 8 Hz) and gamma envelope (0.1 – 16 Hz). We also created low-pass versions of the envelopes using the same rectification procedure but replacing the band-pass filters with a 1^st^ order 0.1 Hz low-pass filter, regardless of carrier frequency. All envelopes and low-pass envelopes were then re-reference to the average of all intracranial channels and then z-scored within each channel. We also kept a second version of the data that was not z-scored. This non-normalized data set was used to extract representative time-courses for brain regions (see below). The envelopes were down-sampled to 100 Hz, and the low-pass envelopes were down-sampled to 10 Hz to reduce memory and CPU demands. The sampling rate of the band-passed data was maintained at 1000 Hz.

#### Functional connectivity

Seed-based functional connectivity maps for a particular data set were computed by calculating the temporal correlation of all channels with that of a specific seed channel or seed region. Similarly, the functional connectomes were computed as the temporal correlation matrix of all channels. Principal components of all connectomes were computed with the *princomp* function in R.

#### Extracting activity of a brain region

To extract activity that is representative of a particular brain region, we first identified all channels in that brain region, and then used the fastICA function in R to identify the time-course of the first principal component. Two slightly different variants of the approach were used. The first version removed the mean across all channels in the region prior to computing the first principal component, the second one did not. The latter approach was used as a default; the former case was used when extracting and comparing multiple regions that may be subject to a shared volume-conducted signal.

#### Intrinsic states

We aimed to identify the emergence and recurrence of stable spatial patterns by computing spatial correlations between different points in time during a resting-state recording. A complete description of the similarity of spatial patterns over time is provided by the spatial correlation matrix. It is related to the temporal correlation matrix that gives rise to the functional connectome, but instead of identifying channels with similar time-courses, it identifies time-points with similar patterns of activity across all channels. In the simplest case, the spatial pattern can be computed from a single data set such as the low-passed delta envelopes. However, it can also be extended to include more than one data set. We had identified the four low-pass envelopes as the most likely substrate for the emergence of intrinsic states. Hence, we concatenated the four data sets along the channel dimension, thus quadrupling the number of channels, and calculated the spatial correlation matrix for this concatenated data set. Two time-points will show strong correlations if they have similar spatial patterns for all four low-passed envelopes. The emergence of stable states would be indicated by high spatial correlations for a contiguous period. A stable state would visually be noticeable as a region of increased width on the main diagonal of the spatial correlation matrix. The recurrence of a stable state should then be visible as a region of increased spatial correlations off of the main diagonal. If the states themselves are stable and the transitions between states is quick, then the regions of increased and decreased spatial correlations will be approximately square-shaped.

### 4.16 Analysis of the delayed frequency discrimination data

As for the other paradigms, data in the delayed frequency discrimination task was re-reference to the average of all intracranial channels. Next, we identified all trials with a lever release, and aligned the data to the onset of the lever movement. We also drew a random sample of the same number of trials without a lever release and aligned them to a simulated lever release time that was randomly drawn from the sample of actual lever release times. We then computed a wavelet transform using the *cwt* function in R. Event-related changes of power as a function of time and frequency were then quantified as the logarithm to the base of 2 of the fractions of the power in trials with vs without a lever response. We identified 15 Hz as carrying the strongest response-related reduction of power. This frequency was used to quantify the temporal dynamics and spatial distribution of response-related beta-band desynchronization. We had also extracted representative time-courses from 7 brain regions (Cerebellum, caudate primary somatosensory cx, primary motor cx, supplementary motor cortex, primary auditory cx, and rostral auditory cx) and used the same procedure to extract the time-course of response-related beta-band desynchronization for these regions rather than individual channels.

## Acknowledgements

The work was supported by MH114223 to TT.

TT, BG, ZA, AH, CMG, MC and KG declare no conflicts of interest. LP and FV are employed by Neuronelektrod, the company that manufactures the T-Probes. NB is employed by Plexon, the company that distributes the T-Probes. The three of them were not involved in the collection, analysis, and interpretation of the data.

