## Supplements for "Volumetric mesoscopic electrophysiology: a new imaging modality for the non-human primate"

### 6. Supplementary Material

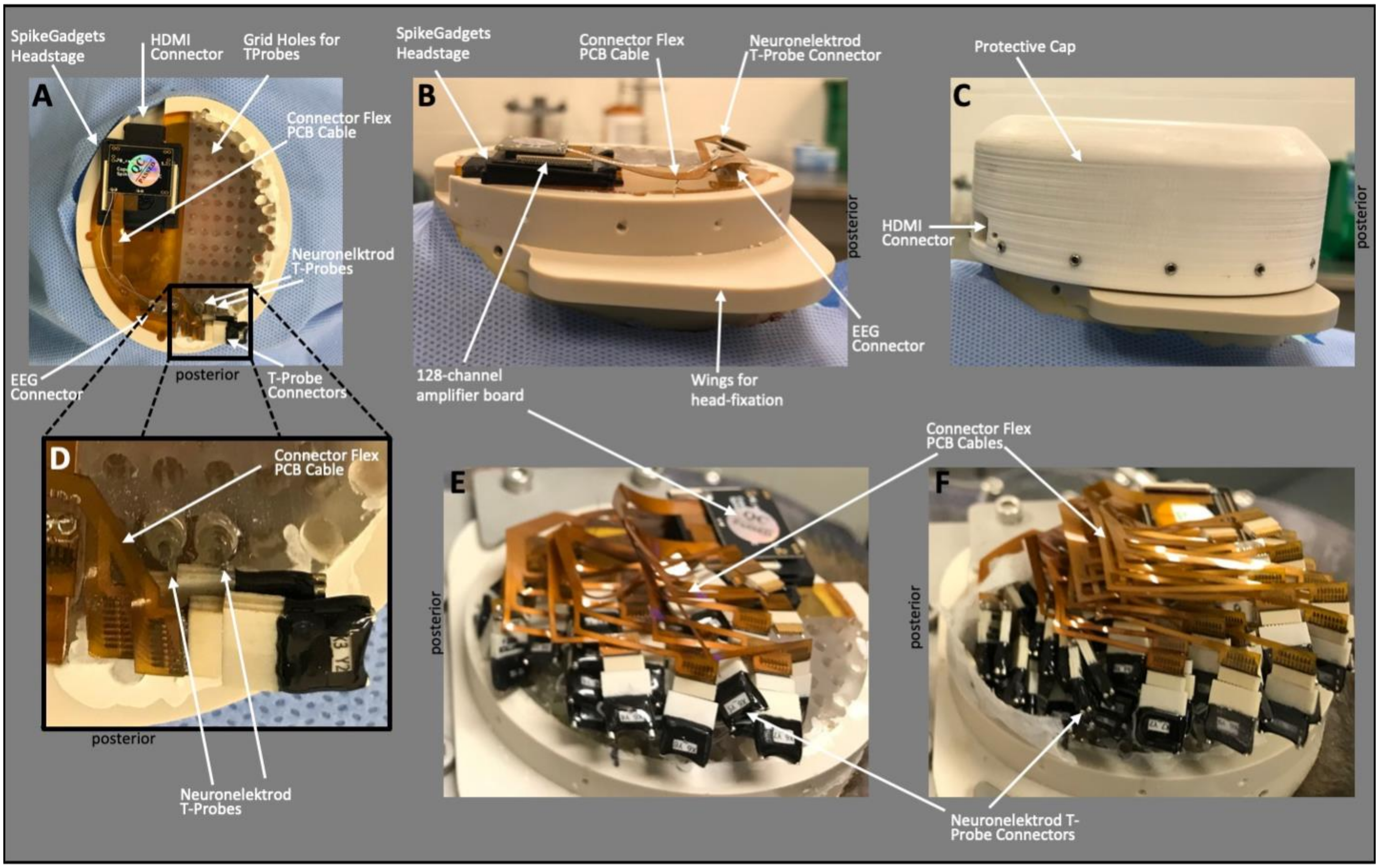


***Supplementary Figure 1 – The MePhys Platform***. *Top-down (**A**) and lateral view (**B&C**) of the MePhys platform on the animal's head. The platform is designed around the 1024 channel modular headstage from SpikeGadgets that is located over the left hemisphere. (**C**) A mini-HDMI connector allows us to stream data without removing the protective cap. Thus, the protective cap can remain in place for months and only needs to be removed for maintenance. (**A**) A regular grid of 72 holes covers the right hemisphere. (**A-D**) The MePhys platform after the first two electrode shafts have been implanted over occipital cortex. The platform after the third (**E**) and final (**F**) surgery is almost completely covered by the Plexon T-Probe connectors and the flex-PCB cables that connect the electrodes to the headstage.*

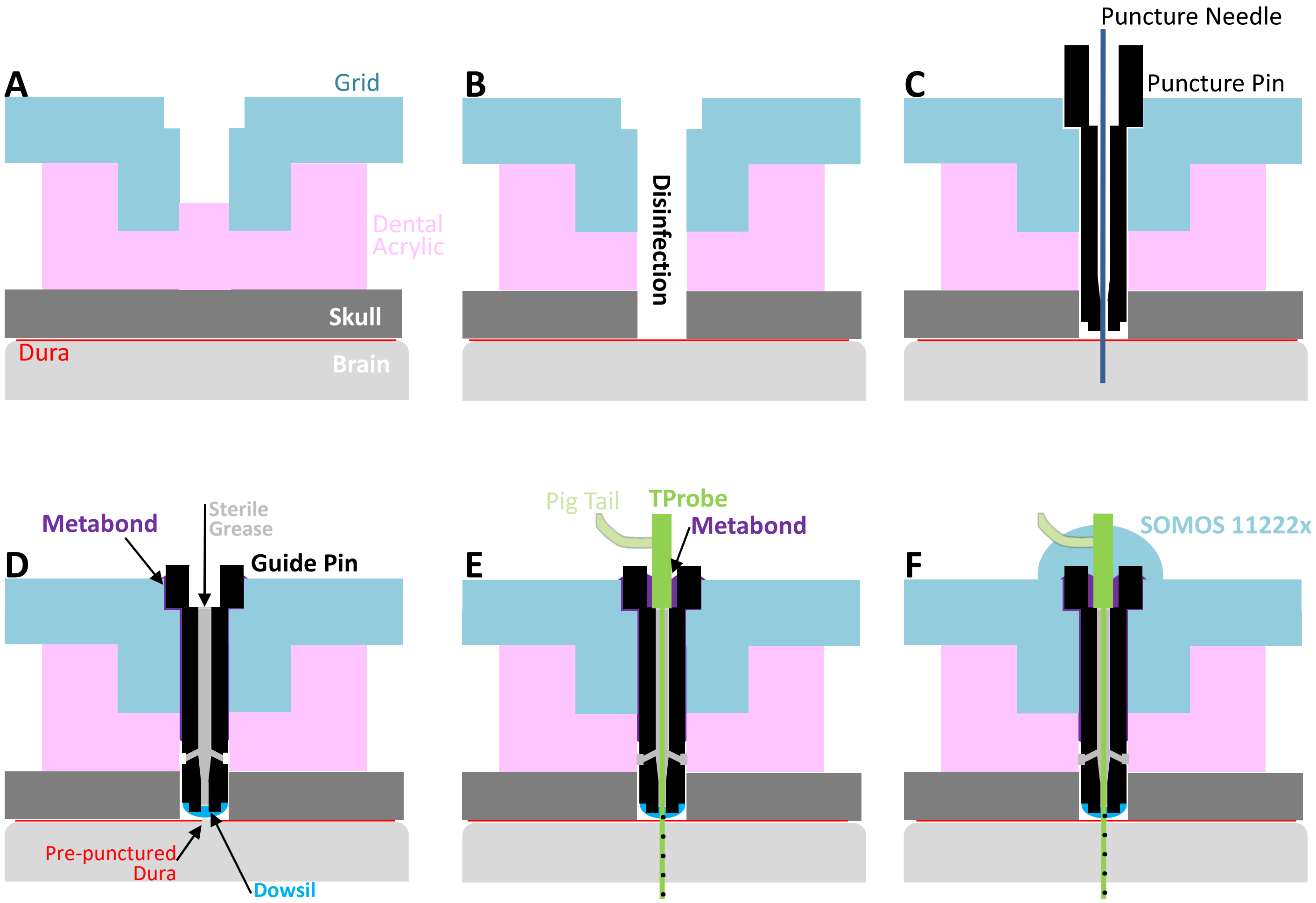


***Supplementary Figure 2 – Electrode implantation****. (**A**) Using custom-designed spacers, a 1.5 mm hole was drilled into the acrylic and skull underlying a specific grid-hole. (**B**) After sterilization, (**C**) a puncture pin was inserted and a needle was used to puncture the underlying dura. (**D**) A sterile grease-filled guide-pin was inserted and cemented into place with metabond. (**E**) The electrode shaft was slowly inserted through the guide pin into the brain by hand and anchored into place with a drop of metabond. (**F**) The top of the electrode was covered with the same 3D printing resin used for the grid itself. The new material was allowed to polymerize in place using a UV lamp.*

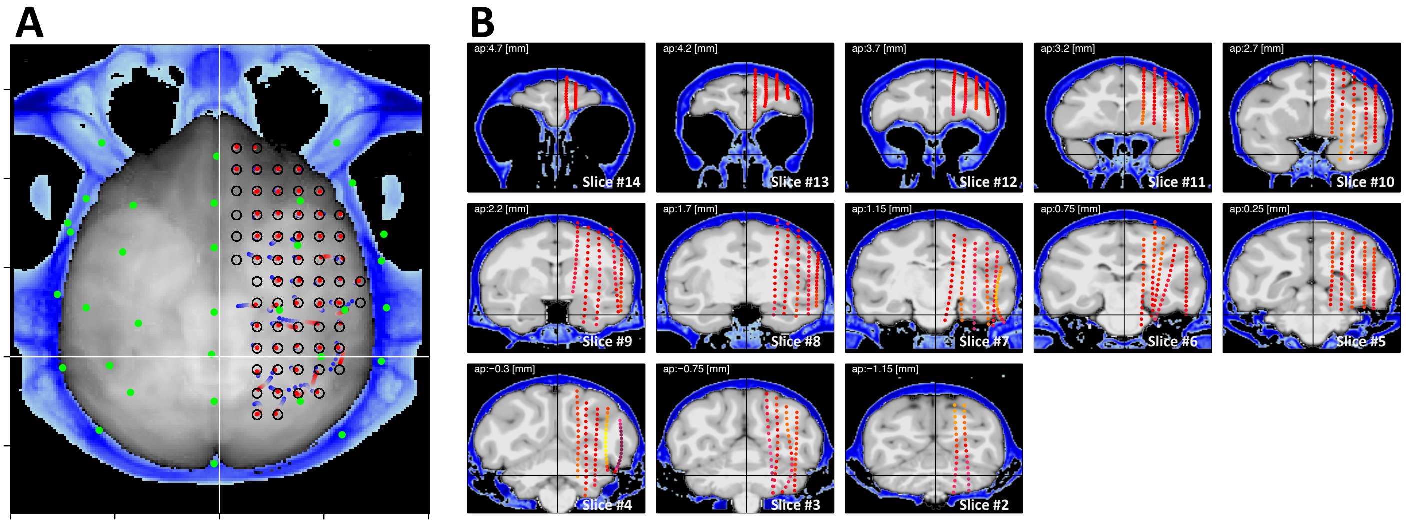


***Supplementary Figure 3 – Electrode locations****. (****A****) EEG (green) and mesoscopic (red to blue) electrode locations in a top-down view. Mesoscopic electrodes at top of the shaft are colored in red, electrodes at the tip in blue. The black circles mark the locations of the grid-holes on the MePhys platform. B) The location of the mesoscopic electrodes in a series of coronal slices. Deviations of electrode position from the coronal slice is indicated by color (yellow: more posterior, red: no deviation, purple: more anterior). Note that most electrode shafts are in the intended location. Four electrode shafts were too long and buckled, two shafts were implanted at an oblique angle.*

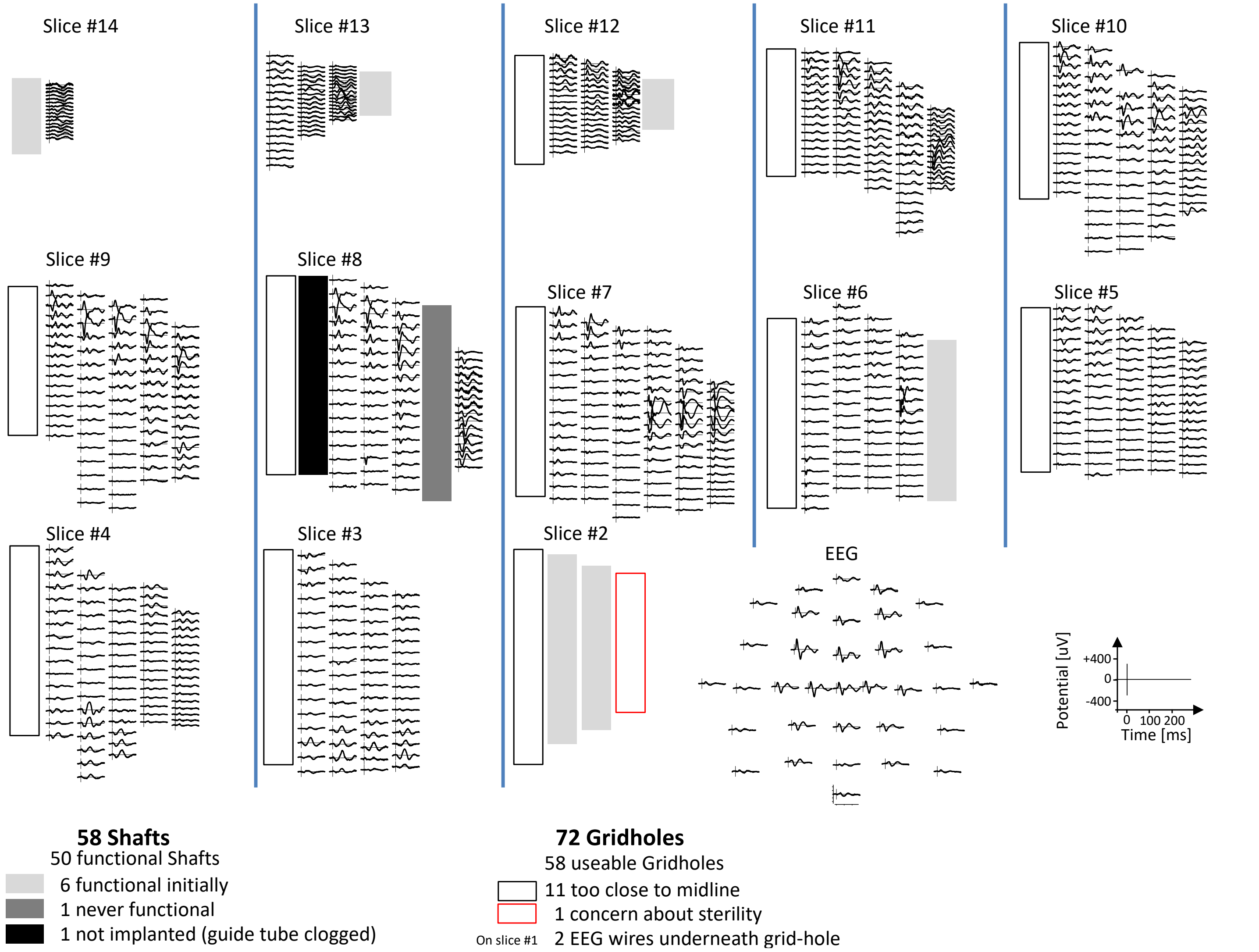


***Supplementary Figure 4 – Meso- and macroscopic auditory evoked potentials**. Auditory evoked responses for intracranial mesoscopic electrodes are arranged in coronal slices. Macroscopic EEG responses are arranged in a top-down view (layout as in Fig 1). The many panels within each slice show the potentials evoked by a brief 85 dB burst of white noise for one electrode contact. For visualization purposes, the evoked potentials at the macroscopic EEG electrodes are multiplied by a factor of 2. The location of the panels approximates the location of the electrode contacts along the 57 functional shafts. Shaded regions indicate the location of electrodes that are not functional. The red and black boxes indicate grid holes that were not considered for implantation. The strongest auditory evoked responses were observed at electrode contacts in the superior temporal plane, motor cortex, cerebellum, prefrontal cortex, and occipital cortex. In addition, we observed auditory evoked potentials at contacts around the medial geniculate nucleus, the inferior colliculus, and the brainstem. These subcortical sources can be appreciated much more clearly using high-pass filter settings.*

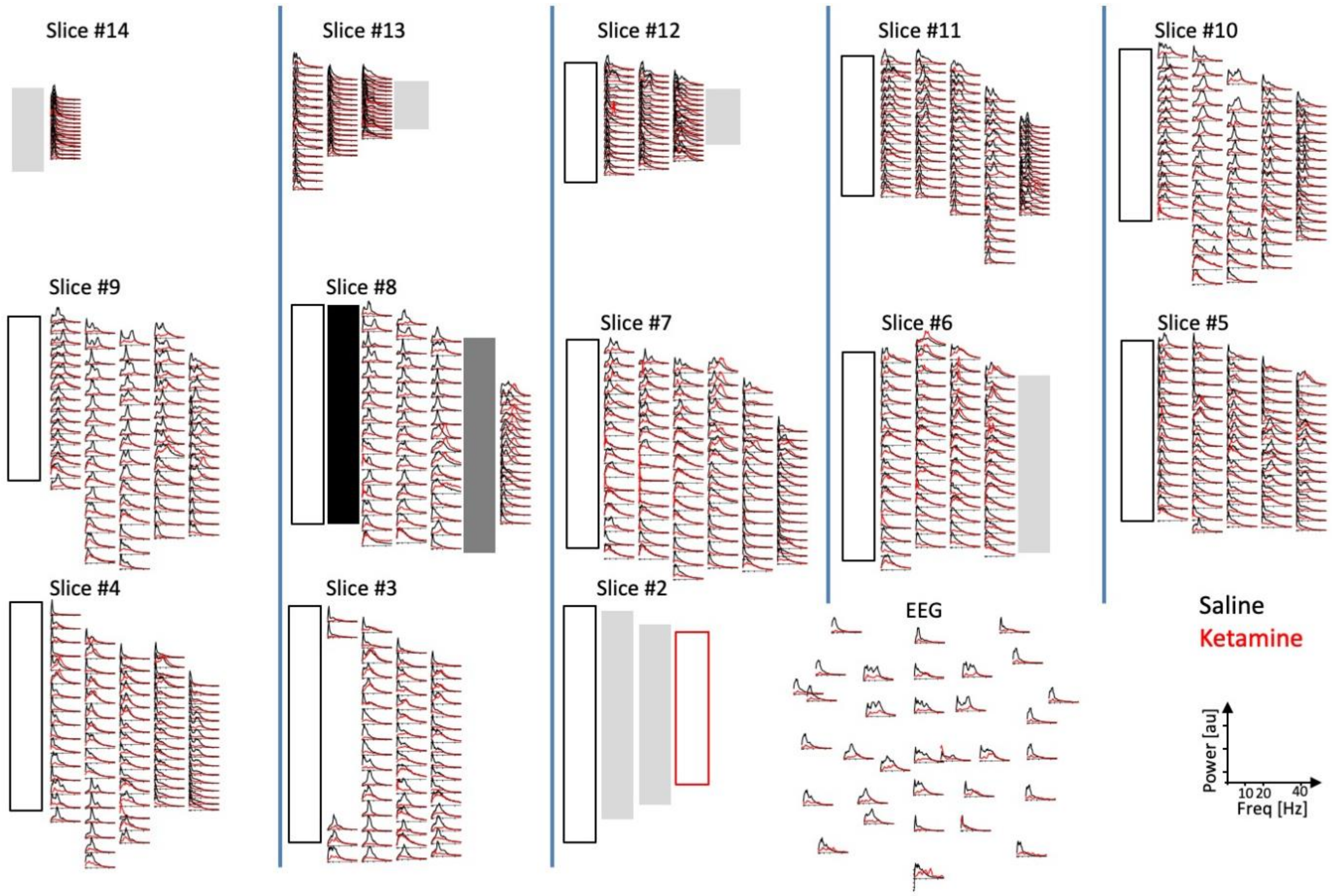


***Supplementary Figure 5 – Resting-state spectra****. Multi-plots of spectra computed in 1-second-long intervals during rest before (black) and after injection of 0.35 mg/kg IM ketamine (red). Layout as in Suppl. Fig 4. Amplitudes in each panel are normalized to a maximum of 1. Ketamine dramatically reduces power and abolishes spectral peaks in most channels.*

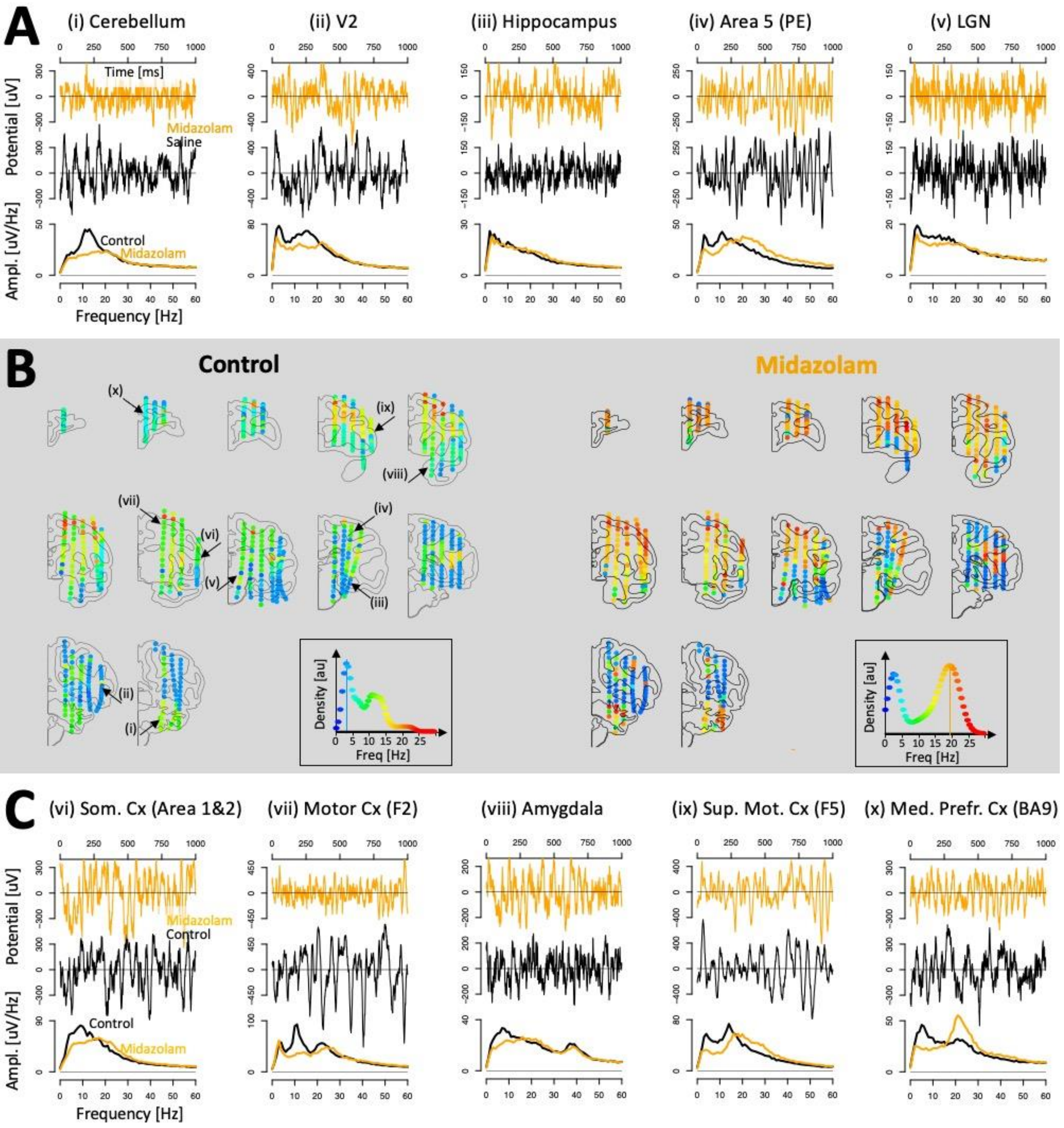


**Supplementary Figure 6 –** ***Distribution of dominant frequencies in the monkey brain.*** *Convention as in Figure 4. Orange lines indicate example traces and spectra following the injection of a subanesthetic dose of midazolam. Note the emergence of beta power across much of prefrontal cortex (e.g., in medial prefrontal cortex; panel x).*

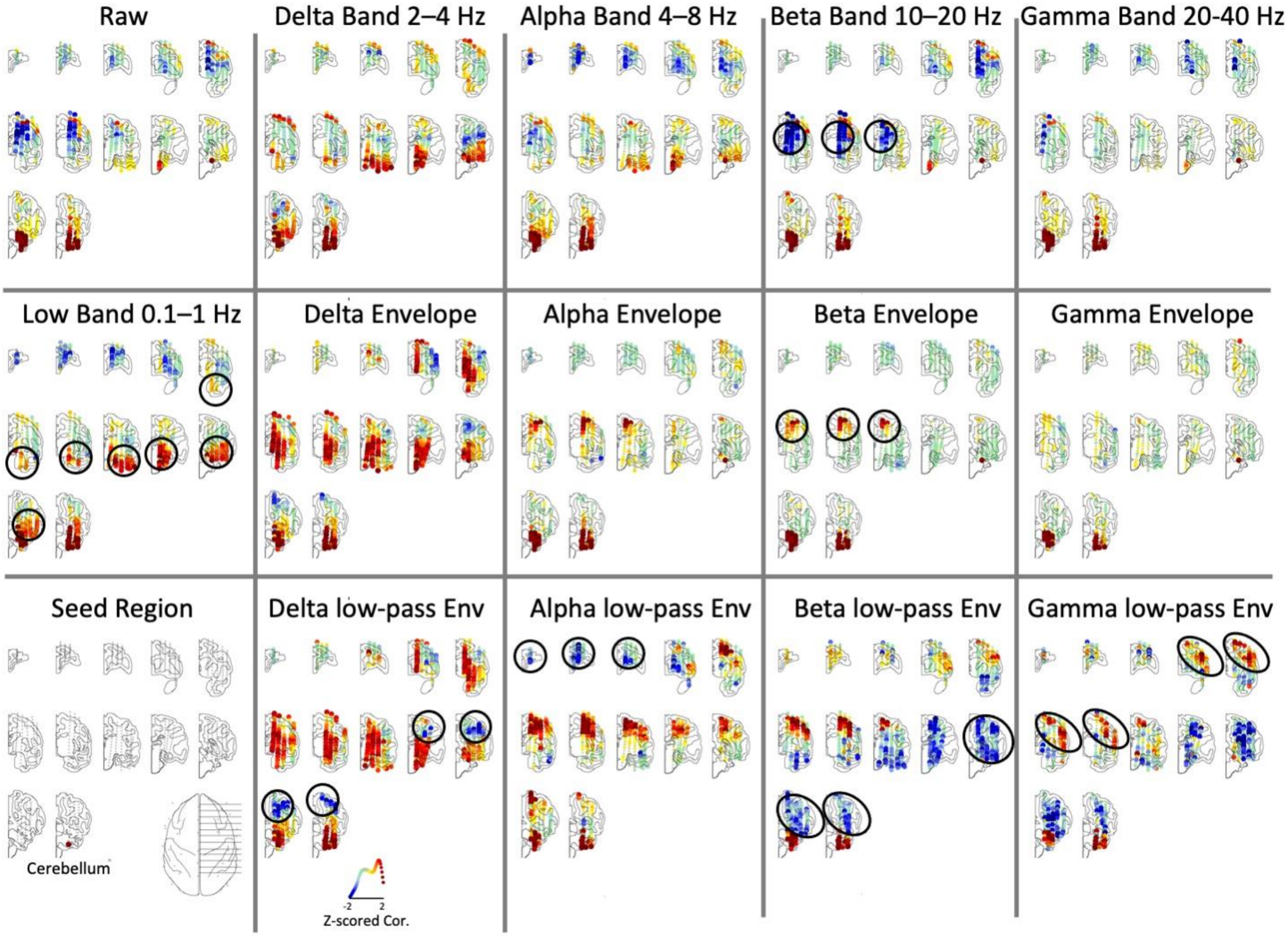


**Supplementary Figure 7 – Functional connectivity depends on carrier frequency and mode**. Functional connectivity maps for the same seed channel in cerebellum (bottom left panel) using 14 different filter settings that use different carrier frequencies (low, delta, alpha, beta, gamma) and modes (band, envelope, and low-passed envelope). We highlighted 7 different spatial connectivity patterns in the map where they are most clearly visible (black ovals). Note, however, that most maps (other than the alpha and beta envelopes) seem to reflect more than just one of the identified spatial patterns. (1) The beta envelope isolated a functional connection from cerebellum to motor cortex and caudate. (2) The gamma low-passed envelopes also identified a connection with motor cortex, but it included more lateral regions of motor cortex and extended into prefrontal cortex. (3) The beta band revealed a functional connection with a broad array of subcortical regions including the basal ganglia and thalamus. (4) The low band highlighted a functional connection to the ventral visual stream and hippocampus. (5) The low-passed alpha envelope identified a connection with the frontal pole. The connectivity maps of the low-pass envelopes are particularly interesting. While most other maps are relatively sparse and identify one or two hot spots, the low-pass envelope maps tend to involve larger contiguous areas, parceling up almost the entire brain into regions of either positive or negative correlation. The parcellation is especially prominent for the delta and alpha low-pass envelopes. In addition, the maps of the low-passed envelopes have overall much stronger correlation values (note that the connection strength here is normalized within each panel, so the absolute correlation strength is not represented).

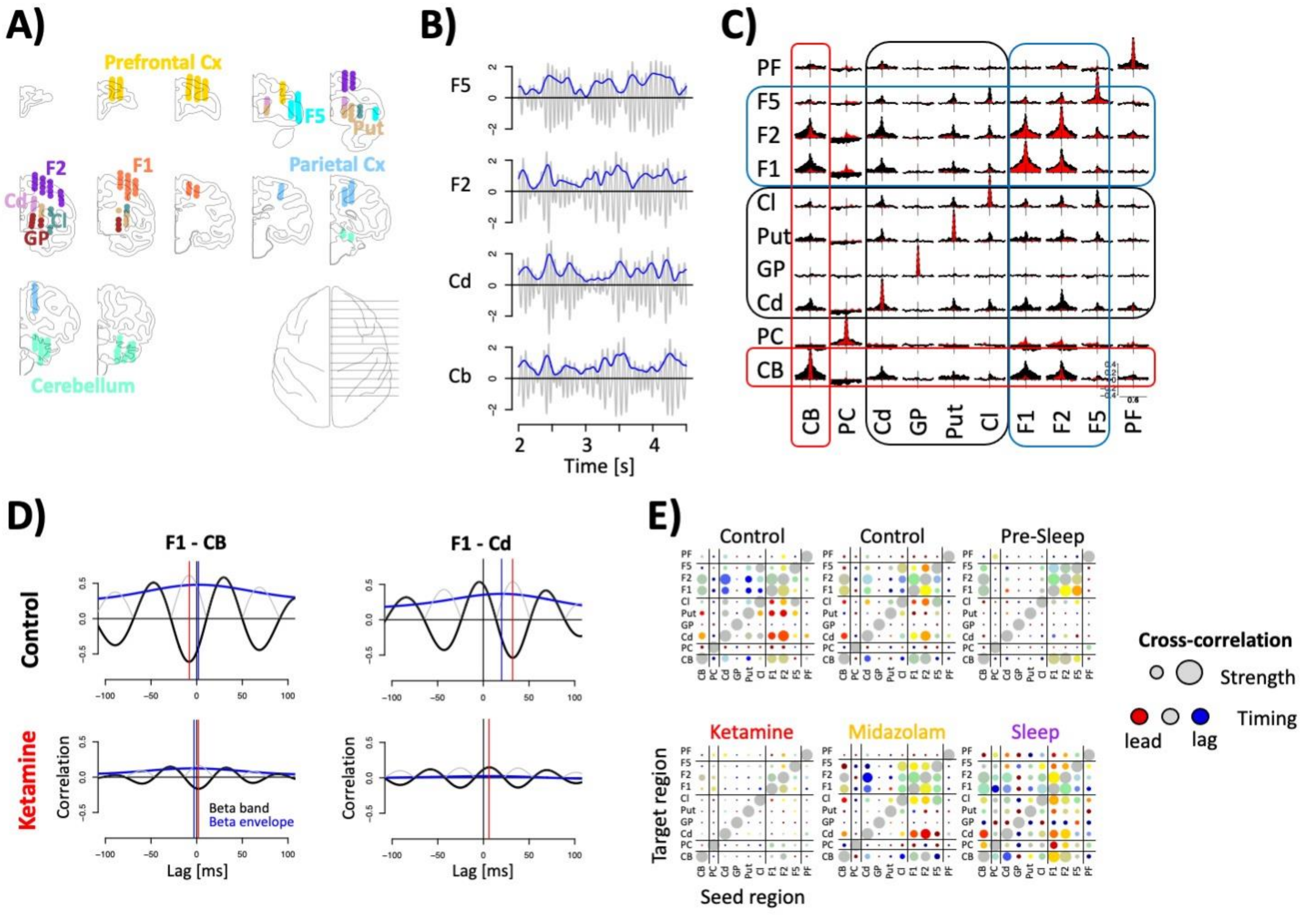


**Supplementary Figure 8 – The beta-band motor network is decoupled by ketamine.** (A) Map of channels in 8 motor and 2 control regions. (B) Example snippets of beta band and beta envelope activity in four example regions. (C) Beta-band cross-correlations during rest between all pairs of regions before (black) and after (red) injection of ketamine. (D) Close-up of beta band (black/gray) and beta envelope (blue) cross-correlations between F1 and CB (left columns) as well as F1 and Cd (right columns), before and after ketamine (top vs bottom). Peaks of the cross correlations are indicated by red (beta band), and blue vertical lines (beta envelope). (E) Cross-correlation strength (dot size) and lag (dot color) for all pairs of regions before (top) and after (bottom) injection of ketamine (left), injection of midazolam (middle) or falling asleep (right).
